# Spatial and Cellular Architecture of the Glioblastoma Microenvironment Associated with Tumor-Infiltrating Lymphocyte Expansion

**DOI:** 10.1101/2025.03.26.645566

**Authors:** Kelly M. Hotchkiss, Kenan Zhang, Anna M. Corcoran, Elizabeth Owens, Jodie Jepson, Kyra Van Batavia, Pamela Noldner, Katayoun Ayasoufi, Kirit Singh, Michael C. Brown, Beth Shaz, Jose Conejo-Garcia, John W. Hickey, Mustafa Khasraw

## Abstract

Tumor-infiltrating lymphocyte (TIL) therapy is effective in several tumor types; however, its feasibility in immune subversive tumors like glioblastoma is unclear. We expanded TILs from glioblastoma specimens and observed marked variability in yield, composition, function, and TCR clonality. By interrogating TILs expanded *ex vivo* alongside their sourced glioma tissue, we sought to identify determinants of successful (TIL^+^) vs unsuccessful TIL expansion (TIL⁻). Expanded TILs were predominantly effector memory CD4⁺ cells and exhibited oligoclonal TCR enrichment. Despite similar T cell abundance in TIL⁺ vs TIL⁻ tumors, TIL⁺ tumors exhibited distinct spatial organization and cellular interactions, including increased endothelial–immune interactions/ proximity and enrichment of vascular-associated niches. CD4⁺ T cells localized near CD68⁺ macrophages in TIL⁺ tumors, while they were positioned near CD163⁺ CD206⁺ macrophages in TIL⁻ tumors. Thus, TIL expansion and functionality are linked to spatial organization and myeloid context; these features may enable biomarker-driven stratification for future TIL therapy in gliomas.

---

Glioblastoma is associated with poor clinical outcomes, with limited advances in systemic therapies over the past two decades^1^. Immunotherapy including immune checkpoint blockade, have not demonstrated clinical benefit in this disease^2–4^, consistent with poor T cells infiltration, myeloid predominance, and multiple mechanisms of immunosuppression. These observations suggest that resistance to immunotherapy in glioblastoma may reflect the organization, functional state, and spatial context of immune cells within the tumor microenvironment (TME)^5,6^.

Adoptive cell therapy using tumor-infiltrating lymphocytes (TILs) is a therapeutic approach that enables the isolation and *ex vivo* expansion of tumor-resident T cells before reinfusion. In other solid tumors like melanoma, this strategy can generate tumor-reactive T cell populations and has been associated with clinical benefit^7,8^. However, its applicability to glioblastoma remains uncertain. Although TIL expansion from glioma has been reported, expanded products vary in efficiency, composition, and function, and the tumor-intrinsic determinants of expansion remain undefined. This variability suggests that TIL expandability may reflect measurable properties of the TME rather than stochastic culture effects.

Studies of glioma have primarily relied on bulk analyses of individual cell populations, which do not capture the spatial organization and intercellular interactions that shape immune function. Emerging single-cell and spatial profiling approaches enable high-resolution characterization of cellular composition, transcriptional states, and spatial relationships. Spatial context has emerged as a critical determinant of anti-tumor immunity, including the positioning of immune cells relative to vasculature, tumor cells, and stromal compartments^9^. In glioblastoma, however, spatial and cellular features associated with effective immune engagement, and with the ability to generate TILs *ex vivo*, are not defined. Clarifying these features has clinical relevance, as it may inform biomarker development, patient selection and guide immunotherapy design.

Here, we integrated ex vivo TIL expansion with multimodal profiling of paired TIL products and source tumors. Expanded TILs were characterized by spectral flow cytometry and T cell receptor sequencing, while tumors were profiled using bulk RNA sequencing, TCR sequencing, single-cell RNA sequencing, in situ spatial transcriptomics, and multiplexed protein imaging. By comparing tumors that generated expandable TILs with those that did not, we identified cellular interactions, spatial organization, and immune states associated with TIL generation in glioblastoma, providing a framework to link TME architecture with therapeutic T cell potential.

## Results

### Patient cohort and definition of TIL expansion status

The study cohort included ten adult patients with IDH wildtype glioblastoma, comprising both primary (n=8) and recurrent (n=3) disease settings (Supplementary Table 1). Tumors were classified based on their ability to generate tumor-infiltrating lymphocytes (‘TIL^+^’ hereafter) using a standardized two-phase expansion protocol developed at our institution (Fig. 1A)^10^. Five TIL⁺ samples met a predefined threshold of ≥10⁸ cells following rapid expansion (REP), a benchmark selected to reflect a clinically meaningful yield compatible with manufacturing and potential therapeutic use, whereas six samples underwent identical processing but did not achieve sufficient cellular outgrowth to meet this threshold (TIL⁻ hereafter).

**Figure 1.**
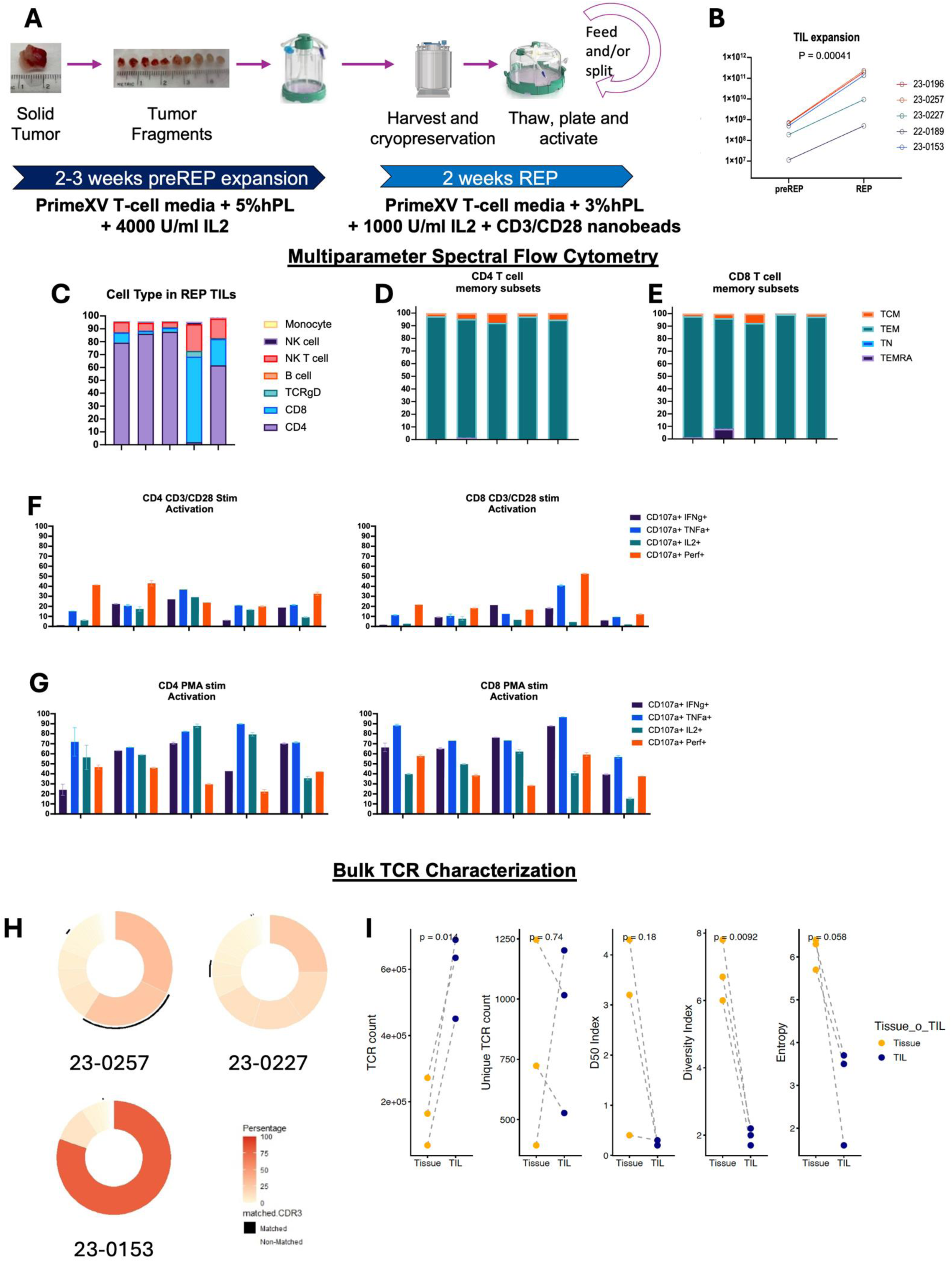
TIL manufacturing, expansion and phenotypic characterization. **A,** TIL manufacturing workflow. Tumor fragments were cultured using a two-phase expansion protocol comprising pre-REP (2–3 weeks; high-dose IL-2) followed by REP (2 weeks; IL-2 plus CD3/CD28 stimulation), with intermediate cryopreservation and reactivation steps. **B,** TIL expanded robustly from pre-REP to REP across all paired samples, despite substantial inter-patient variability in expansion magnitude from an equivalent starting tumor mass of 0.5 g. Because cell yields spanned several orders of magnitude, paired analysis was performed on log10-transformed cell counts, demonstrating a significant increase after REP (n = 5, paired t-test, P = 0.00041; mean expansion, 142-fold). The y-axis shows log10-transformed cell counts. **C,** Cellular composition of REP TIL products. REP-expanded TILs were predominantly CD4⁺ T cells in most samples, with one case (23-0153) enriched for CD8⁺ T cells. Minor populations included NK, NKT, γδ T cells and B cells. **D – E,** Memory phenotype of REP TILs. Both CD4⁺ and CD8⁺ TIL populations were composed predominantly of effector memory (TEM) cells (>90%), with minimal naïve (TN), central memory (TCM) or TEMRA subsets. **F–G,** Functional activation capacity. PMA stimulation induced cytokine production in a substantial fraction of CD4⁺ and CD8⁺ TILs (∼70–90%). CD3/CD28 stimulation elicited lower responses (∼20–40% in CD4⁺ and ∼15–50% in CD8⁺ cells), indicating variable responsiveness to physiologic TCR signaling. **H–I,** TCR repertoire characterization. TCR sequencing revealed oligoclonal expansion within REP TIL products, with variability in clonotype distribution, diversity and entropy across patients, consistent with selective outgrowth of dominant clones during *ex vivo* expansion.

### TIL expansion from glioma is feasible but yields heterogeneous products

Samples were classified as TIL⁺ if they achieved ≥10⁸ cells following REP (n=5) and were carried forward for phenotypic and functional analyses. In contrast, products from tumors classified as TIL⁻ (n = 6) underwent the same culture protocol but failed to achieve productive expansion, yielding <10⁸ cells. Because these cultures generated too few cells for phenotypic analysis or functional characterization, they were included as non-expanding controls.

TIL⁺ samples showed robust proliferation from pre-REP to REP, with increased cell yield observed in every paired sample despite equivalent starting tumor input of 0.5 g (Fig. 1B). Because cell yields spanned several orders of magnitude, paired analysis was performed on log10-transformed counts, confirming a significant increase after REP (n = 5, paired t-test, P = 0.00041; geometric mean expansion, 142-fold). The magnitude of expansion varied across patients, indicating marked inter-patient differences in proliferative capacity under standardized culture conditions.

Phenotypic characterization of REP products by spectral flow cytometry demonstrated that expanded TILs were predominantly CD4⁺ T cells in most samples, with one case (23-0153) showing relative CD8⁺ enrichment (Fig. 1C; Supplementary Fig. 1A–C). While lineage dominance (CD4⁺ versus CD8⁺) varied across cultures, overall immune cell composition was largely preserved between pre-REP and REP phases, indicating that expansion did not alter major lineage distributions (Fig. 1C; Supplementary Fig. 1C).

Both CD4⁺ and CD8⁺ compartments were overwhelmingly composed of effector memory (T_EM) cells (>90%), with minimal representation of naïve (T_N), central memory (T_CM), or Terminally Differentiated Effector Memory cells re-expressing CD45RA (TEMRA) subsets (Fig. 1D–E). Functional profiling indicated that expanded TILs retained substantial effector capacity, as evidenced by robust cytokine production following pharmacologic stimulation, with approximately 70–90% of cells producing cytokines in response to PMA (Fig. 1G). In contrast, stimulation through CD3/CD28 elicited lower and more variable responses (approximately 20–50%), implying heterogeneity in TCR-dependent activation potential across samples (Fig. 1F; Fig. 1G; Supplementary Fig. 1E). Activation markers were detectable in both CD4⁺ and CD8⁺ compartments, whereas markers associated with suppression or exhaustion were generally low (Supplementary Fig. 1A–B). These data suggest that expanded TIL populations are not globally dysfunctional but instead comprise antigen-experienced T cells with preserved effector capacity and variable sensitivity to receptor-mediated stimulation, consistent with differences in activation state rather than uniform exhaustion.

TCR sequencing revealed oligoclonal expansion within REP products, characterized by reduced diversity and dominance of select clonotypes (Fig. 1H–I). Tracking of shared clonotypes between tumor and matched TIL products demonstrated partial overlap, with preferential expansion of specific subsets of tumor-derived clones during *ex vivo* culture (Supplementary Fig. 1F), potentially implying selective outgrowth of tumor-reactive populations. In addition to αβ T cells, expanded TIL products contained variable proportions of innate-like lymphocyte populations, including NKT and γδ T cells. Variability in TCRγδ subset composition (γδ1 versus γδ2) (Supplementary Fig. 1D) implied patient-specific differences in innate-like T cell contributions. Several γδ TCR clones were detected by bulk TCR sequencing (Supplementary Fig. 1H). These data demonstrate substantial heterogeneity in lineage composition, functional responsiveness, and clonal architecture across expanded TIL products despite a standardized manufacturing process.

### Cellular composition across TIL□ and TIL□ tumors

We next compared tumors that generated sufficient *ex vivo* expanded TIL products for downstream characterization (TIL⁺, n=5) with tumors that failed to yield adequate outgrowth (≥10⁸ cells) under identical culture conditions (TIL⁻, n=6). We used complementary single-cell RNA sequencing (scRNA-seq), in situ spatial transcriptomics (Xenium, custom panel), and CO-Detection by indEXing (CODEX; PhenoCycler), an ultra-high-plex immunofluorescence platform enabling spatial single-cell imaging of >40 selected protein markers in one tissue section. Xenium and CODEX were performed on the original tumor tissues, providing orthogonal yet convergent views of the TME.

Across modalities, UMAP projections consistently resolved major cellular compartments, including malignant, lymphoid, myeloid, and stromal populations, with substantial overlap in their global distributions between TIL⁺ and TIL⁻ tumors (Fig. 2A–B). Each platform provided distinct resolution of cellular features: scRNA-seq defined transcriptionally distinct tumor cell states (MES-like, AC-like, NPC-like, OPC-like) and immune subsets; Xenium preserved spatially resolved transcriptomic identities within intact tissue architecture; and CODEX enabled protein-level phenotyping of immune and stromal populations, including macrophage polarization states and lymphocyte subsets defined by surface marker expression.

**Figure 2.**
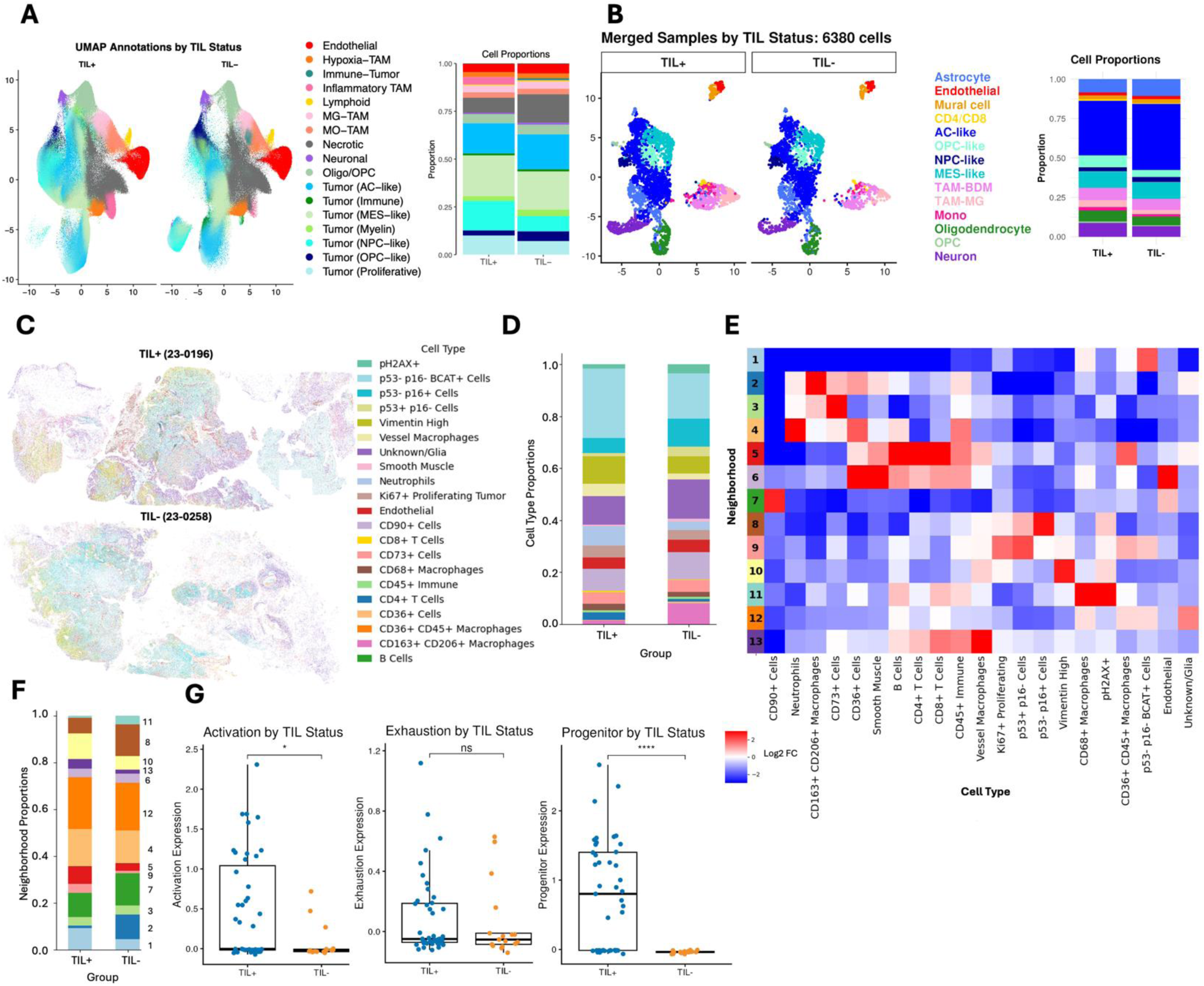
Cellular composition, spatial organization, and lymphoid cell states according to TIL expansion status. A, Xenium UMAP visualization. UMAP projections of Xenium spatial transcriptomic profiles annotated by major cell types, shown separately for TIL⁺ and TIL⁻ tumors. Adjacent bar plots summarize relative cell-type proportions. B, scRNA-seq UMAP visualization. Integrated single-cell RNA-seq data (n = 6,380 cells) displayed as UMAPs stratified by TIL status with corresponding lineage proportions. C, Spatial distribution of cell types. Representative CODEX spatial maps illustrating regional organization of tumor and immune compartments. D, Cell-type composition by group. Stacked bar plots summarizing annotated cell populations across TIL⁺ and TIL⁻ tumors. E, Neighborhood analysis. Heatmap of neighborhood enrichment scores (log fold change) identifying differences in local cell–cell organization. F, Neighborhood composition. Relative abundance of inferred spatial neighborhoods across groups. G, Lymphoid cell-state scores. Boxplots of lymphoid gene-signature scores, including activation, exhaustion and progenitor-associated programs, compared between TIL⁺ and TIL⁻ tumors.

Cell type annotation across modalities relied on multiple complementary markers. scRNA-seq leveraged canonical transcriptional programs (e.g., SOX2, OLIG2 for tumor lineage; CD3D, CD4, CD8A for T cells; CD68, CX3CR1 for myeloid populations), Xenium applied targeted spatial gene panels for simultaneous localization and classification, and CODEX used antibody-based detection of lineage and functional markers (e.g., CD3, CD4, CD8, CD68, CD163, CD206) to resolve immune phenotypes *in situ*. This multimodal approach enabled both cross-validation of shared cell classes and extension into modality-specific domains.

Although tissue organization showed marked intra- and inter-sample heterogeneity (Fig. 2C), CODEX-based quantification revealed no consistent group-level differences in cellular composition between TIL⁺ and TIL⁻ tumors. Individual cell-type proportions varied substantially across samples without reproducible enrichment of any population, including T cells, by TIL status (Fig. 2D; Supplementary Fig. 2B, D, F). Similar results were observed after collapsing annotations into broader lineages, where apparent differences were largely driven by sample-specific variation. After ontology harmonization, lineage-level proportions were concordant across scRNA-seq, Xenium, and CODEX within matched samples, arguing against platform-specific bias. Given the marked tumor heterogeneity and limited cohort size, grouped comparisons should be interpreted cautiously.

To investigate hierarchical cellular organization within these tumors, we performed neighborhood analysis on our CODEX data^11^. We defined windows of 20 nearest neighbors around each cell and clustered them using k-means to identify 13 distinct neighborhoods (Fig. 2E). We then compared neighborhood abundances between TIL⁺ and TIL⁻ tumors and, like our cell type–level analysis, observed no significant enrichment of any neighborhood in either group. Comparison of lymphoid cell states between TIL⁺ and TIL⁻ tumors (Fig. 2F and 2G) revealed significantly increased expression of gene signatures associated with activation (p<0.05) and progenitor (p<0.00001) programs, with non-significant increases in exhaustion status also observed.

These data indicate that cellular composition, whether defined by transcriptional states, spatial cell types, or protein phenotypes, is heterogeneous across glioblastoma and does not clearly segregate by TIL expansion capacity in this cohort. Although limited by sample size, inter-patient variability, and lack of normalization to baseline T cell input, these findings suggest that bulk differences in major cellular populations, including TIL levels, may not explain TIL expandability, supporting further investigation of spatial relationships, microenvironmental context, and intercellular interactions.

### Endothelial–immune interactions and spatial organization associate with TIL expansion

We next examined whether spatial organization, defined by cellular positioning, proximity relationships, and neighborhood architecture within the TME, distinguishes tumors capable of generating TILs.

Neighborhood analysis defined a discrete microenvironmental niche, Niche 3, characterized by coordinated enrichment of endothelial cells, lymphoid populations, and monocyte-derived tumor-associated macrophages (Fig. 3A). This niche was preferentially represented in TIL⁺ tumors (Fig. 3B), suggesting an association between vascular-associated immune organization and TIL expandability. Transcriptional features within this niche included extracellular matrix components (DCN, COL1A2, THBS1), myeloid-associated markers (CD163, CD68, APOE), and immune-regulatory molecules (HLA-E, TGFBI), implying a myeloid-influenced TME. Analysis of lymphoid cell niche distribution showed a higher proportion of lymphoid cells within Niche 3 in TIL⁺ tumors, consistent with an association between endothelial–lymphoid interactions and successful TIL expansion (Supplementary Fig. 3A). This was supported by direct visualization, which demonstrated clustering of lymphoid cells along vascular structures in TIL⁺ tumors, whereas lymphoid cells in TIL⁻ tumors were more diffusely distributed and less consistently associated with vasculature (Fig. 3C). Among the six spatial neighborhoods identified, Niche 1, characterized by enrichment for necrotic neighboring cells, contained a significantly higher proportion of lymphoid cells in TIL⁻ tumors compared with TIL⁺ tumors (p < 0.05; Supplementary Fig. 3A). These findings indicate that lymphoid cell presence alone did not predict successful TIL expansion. Instead, lymphoid cells in non-expanding tumors were more frequently localized within necrosis-associated contexts that may be less supportive of productive T cell survival, activation, or recovery, highlighting the importance of tissue organization in relation to TIL expandability.

**Figure 3.**
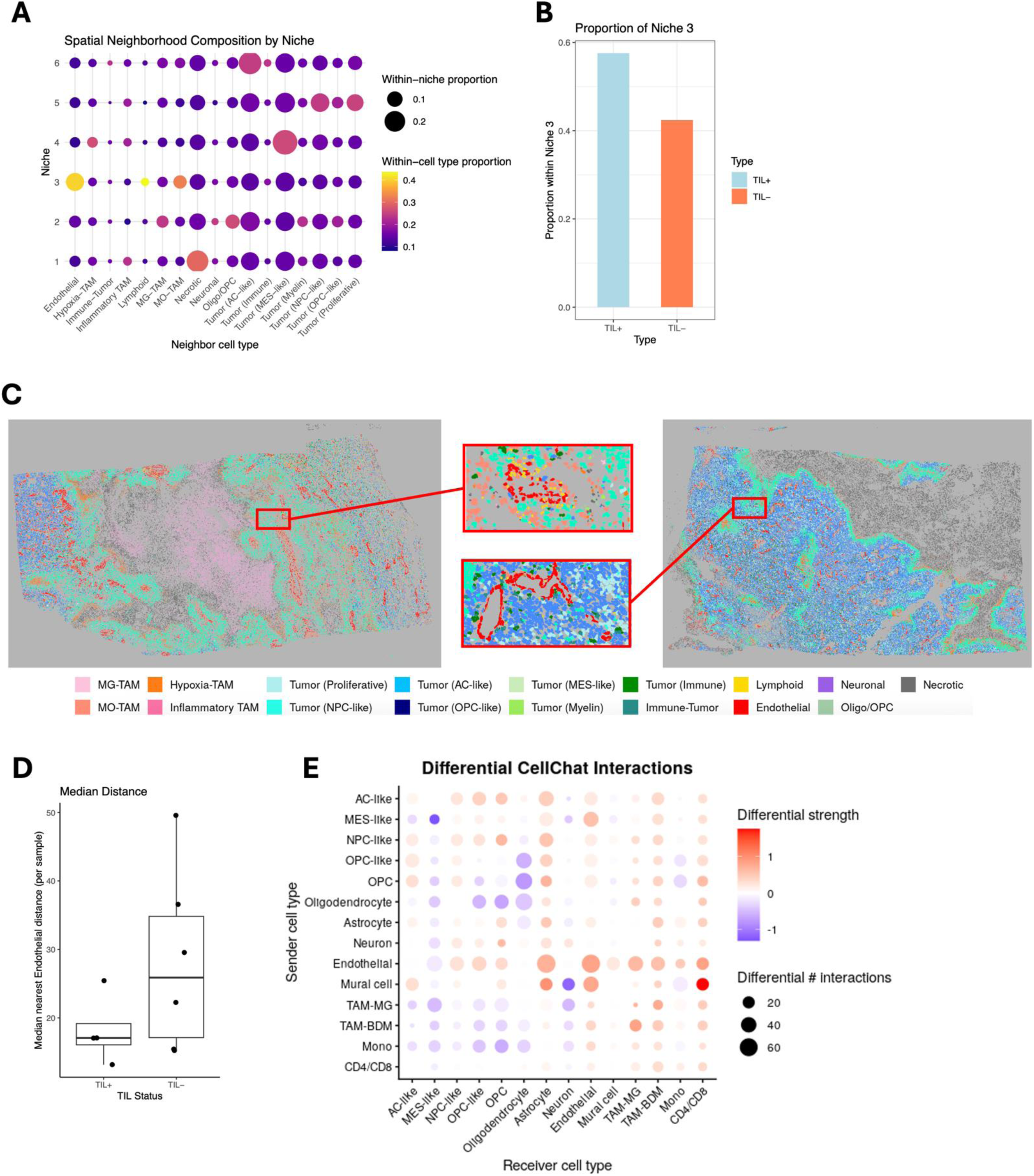
Spatial endothelial–immune organization and multicellular communication associated with TIL expandability in glioblastoma. A, Spatial neighborhood composition. Dot size indicates within-niche cell-type proportion and color intensity indicates contribution of each cell type to the niche. A niche enriched for endothelial, lymphoid and monocyte/macrophage populations was identified (Niche 3). B, Relative abundance of Niche 3. Niche 3 was more abundant in TIL⁺ tumors. C, Representative spatial maps. Spatial maps from TIL⁺ and TIL⁻ tumors with insets highlighting endothelial–immune interactions. D, Distance to endothelial cells. Median nearest distance to endothelial cells across tumors stratified by TIL status. Boxes indicate interquartile range, center lines indicate medians and whiskers indicate range. E, Differential cell–cell communication. Heatmaps showing differences in inferred interaction number and strength between TIL⁺ and TIL⁻ tumors. Bar plots summarize total outgoing and incoming interactions by cell type.

Also, there was a trend toward endothelial-associated localization of lymphoid cells in TIL⁺ tumors (Fig. 3D, p = 0.33, Wilcoxon test). Although not statistically significant, lymphoid cells tended to be closer to endothelial structures, with reduced median lymphoid–endothelial distances in TIL⁺ relative to TIL⁻ tumors.

Analysis across endothelial contour gradients showed that lymphoid cell distribution varied as a function of vascular proximity rather than being uniformly dispersed, with this spatial pattern observed across tumors irrespective of TIL expansion status (Supplementary Fig. 3B–C), indicating that immune localization is spatially constrained and context-dependent. Lymphoid cells in a perivascular neighbor showed significantly higher progenitor and lower exhaustion scores compared to non-perivascular lymphoid cells (p<0.01, Wilcoxon test; Supplementary Fig. 3D).

CellChat analysis of scRNA-seq data further supported these findings by inferring major signaling inputs, outputs, and network-based intercellular communication^12^. Consistent with spatial analyses, CellChat revealed greater density and strength of predicted intercellular interactions centered on endothelial populations in TIL⁺ tumors. Endothelial cells showed more frequent interactions with CD4⁺ and CD8⁺ T cells, tumor cells, astrocytes, and mural cells, while CD4⁺ and CD8⁺ T cells showed stronger interactions with endothelial and mural compartments (Fig. 3E). This suggest that differences associated with TIL expandability are driven less by overall cellular abundance and more by the spatial organization and signaling architecture of the TME.

### CXCL12-CXCR4 signaling shapes lymphoid cell state in TIL samples

To better understand how spatial location influences different lymphoid cell states between TIL^+^ and TIL⁻ groups, we performed unbiased spatial variant gene and spatial cell communication analyses. BANKSY^13^, a spatial transcriptomic method that incorporates neighborhood gene expression to define each cell’s local molecular environment, was used to quantify environmental gene expression surrounding lymphoid cells. The volcano plot identified significantly different genes across the TIL^+^ and TIL⁻ groups(Supplementary Fig. 4A), including multiple chemokines, cytokines, and immune-related factors, indicating that several signaling pathways may contribute to these differences. Gene set enrichment analysis (GSEA), which tests whether predefined gene sets differ between biological states^14,15^, revealed broadly concordant immune-related programs in TIL⁺ and TIL⁻ tumors (Supplementary Fig. 4B). We therefore examined cytokine–receptor expression and inferred intercellular signaling across cell types. Among the detected interactions, CXCL12 was enriched in endothelial cells, whereas its receptor CXCR4 was enriched in lymphoid cells (Supplementary Fig. 4C). Spatial CellChat analysis showed stronger CXCL12–CXCR4 signaling received by lymphoid cells in TIL⁺ tumors, a finding supported by the scRNA-seq analysis (Supplementary Fig. 4D, E). Although other lymphoid-trafficking pathways, including CCL19–CCR7, were also considered, CellChat did not identify these interactions as significant in either TIL⁺ or TIL⁻ tumors. This may reflect weak or context-dependent signaling, limited T cell representation, or filtering thresholds inherent to ligand–receptor inference algorithms.

In addition, the interactome network showed within-lymphoid self-signaling, suggesting communication within the pooled lymphoid compartment, although combined annotation precluded definitive assignment of CD4–CD8 directional interactions (Supplementary Fig. 4D, E). A trend toward a higher proportion of CXCR4⁺ lymphoid cells was observed in TIL⁺ tumors, but this was not significant (Supplementary Fig. 4F, Wilcoxon test, p = 0.2694). Spatial mapping showed heterogeneous CXCL12 and CXCR4 expression across tumors and regions, without consistent enrichment in either TIL⁺ or TIL⁻ groups (Supplementary Fig. 4G, H).

Thus, chemokine signaling differences between TIL⁺ and TIL⁻ tumors were less apparent than differences in spatial organization, including vascular-associated immune niches marked by coordinated endothelial–lymphoid–myeloid interactions and increased vascular-centered communication. These niches may identify T cells retained in perivascular or less tumor-infiltrated regions and therefore relatively spared from terminal dysfunction or deletion.

### Macrophage-associated spatial organization and functional states differ between TIL⁺ and TIL⁻ tumors

CODEX imaging resolved lymphoid- and macrophage-associated spatial context at the protein level (Fig. 5). TIL⁺ tumors showed a trend toward increased CD4⁺ T cell abundance, whereas TIL⁻ tumors showed a trend toward increased CD163⁺CD206⁺ macrophages (Fig. 4A, B). Although overall cell-type composition was heterogeneous, CD163⁺CD206⁺ macrophages were positioned closer to T cells in TIL⁻ tumors (Fig. 4C, D). These patterns were not observed for CD8⁺ T cells or CD68⁺ macrophages, suggesting that they reflect distinct immune programs rather than broad lymphoid or myeloid shifts (Supplementary Fig. 5A, B). Consistent with this, a CD163⁺CD206⁺ macrophage-enriched neighborhood trended toward enrichment in TIL⁻ tumors, whereas a lymphoid-enriched neighborhood containing CD4⁺ and CD8⁺ T cells trended toward enrichment in TIL⁺ tumors, although neither reached statistical significance (Fig. 4E, F).

**Figure 4.**
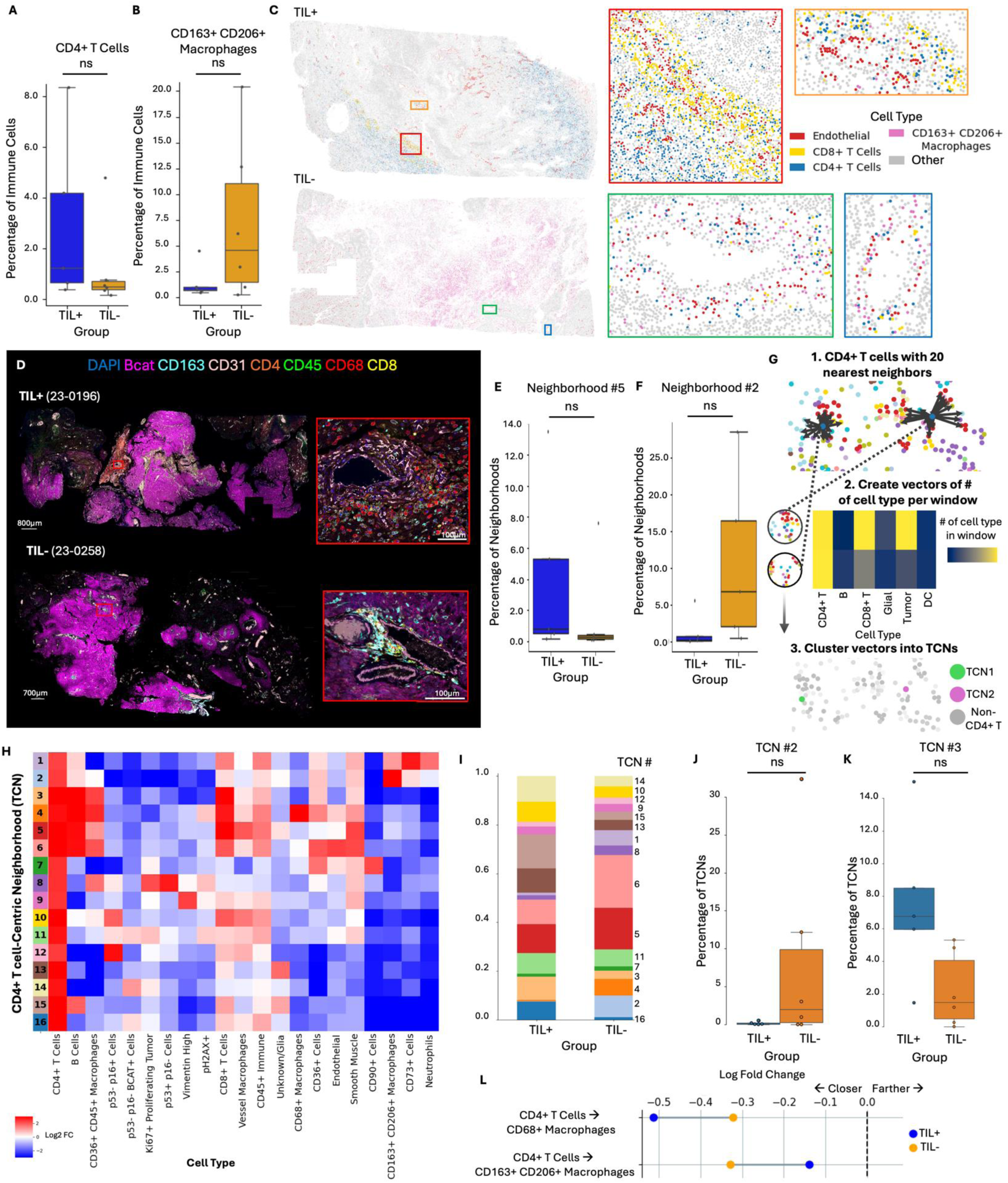
CODEX single-cell spatial proteomics reveals macrophage-associated organization in TIL⁺ and TIL⁻ tumors. A, CD4⁺ T-cell abundance. Proportion of CD4⁺ T cells among immune cells across groups. B, CD163⁺CD206⁺ macrophage abundance. Proportion of CD163⁺CD206⁺ macrophages among immune cells across groups. C, Spatial localization of selected cell types. Representative overlays showing CD4⁺ T cells, CD8⁺ T cells, endothelial cells and CD163⁺CD206⁺ macrophages. D, Representative CODEX images. Images of TIL⁺ and TIL⁻ tumors highlighting immune, endothelial and tumor populations. E, Lymphoid-enriched neighborhood abundance. Relative abundance of Neighborhood 5 across groups. F, Macrophage-enriched neighborhood abundance. Relative abundance of Neighborhood 2 across groups. G, Cell-centric neighborhood analysis schematic. Overview of neighborhood analysis workflow. H, CD4⁺ T-cell-centric neighborhood heatmap. Heatmap showing cell-type abundance within neighborhoods derived from the 20 nearest neighbors around CD4⁺ T cells. I, Neighborhood composition. Relative abundance of T-cell-centric neighborhoods across groups. J, CD8⁺ T-cell-enriched neighborhood abundance. Relative abundance of CD8⁺ T-cell-enriched TCN3 across groups. K, Macrophage-enriched neighborhood abundance. Relative abundance of macrophage-enriched TCN2 across groups. L, Cell–cell interaction analysis. Permutation-based analysis showing non-random proximity of CD4⁺ T cells to macrophage populations.

**Figure 5.**
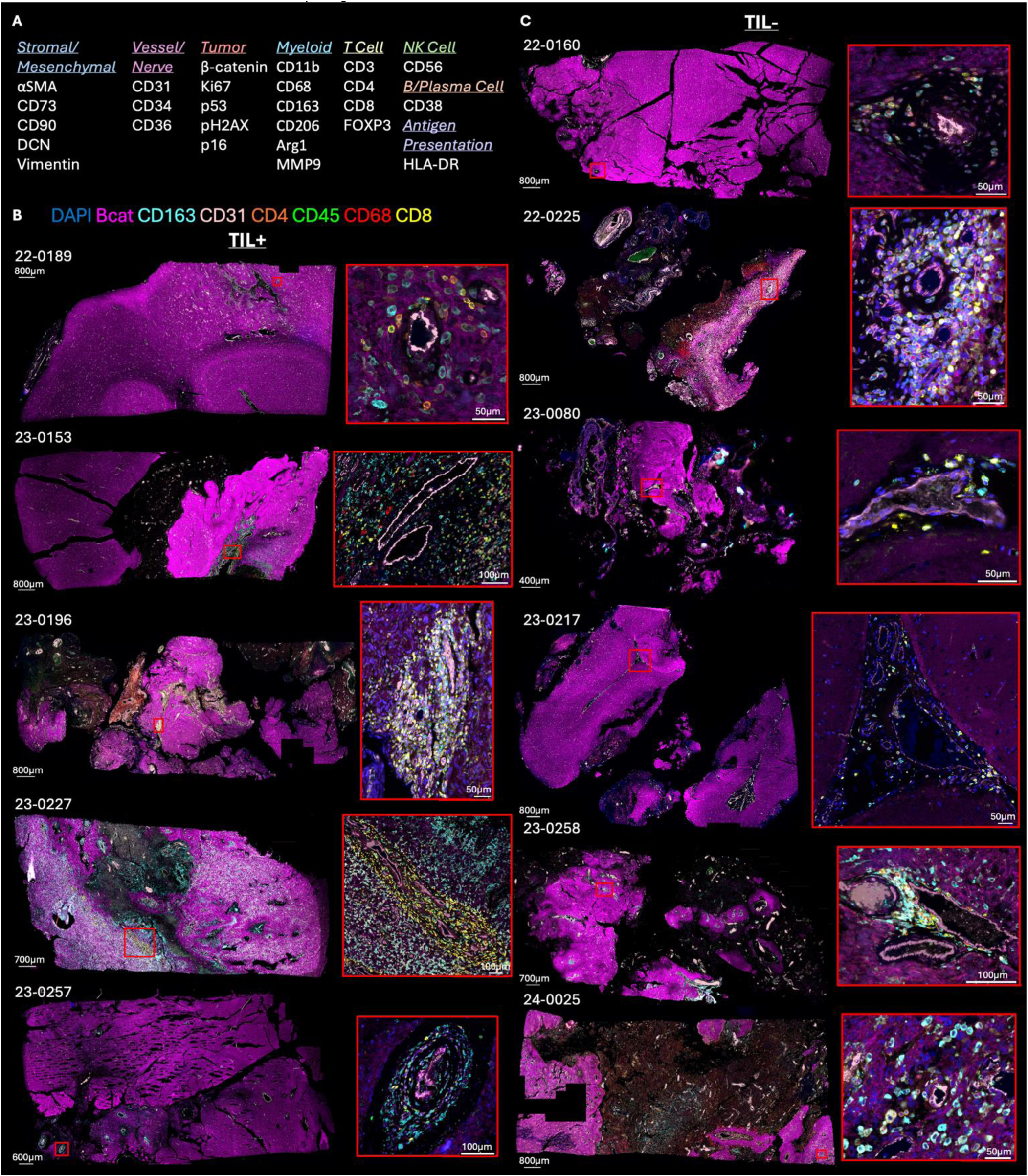
CODEX multiplexed imaging reveals immune–vascular architecture according to TIL expansion status. A, Marker panel. CODEX marker panel used to annotate tumor, stromal, endothelial, myeloid and lymphoid compartments. B, TIL⁺ tumors. Representative glioblastoma samples with successful TIL expansion. Insets highlight immune infiltration, T cells and vasculature. C, TIL⁻ tumors. Representative glioblastoma samples without successful TIL expansion. Insets highlight immune infiltration, T cells, CD163⁺ macrophages and vasculature.

Given the predominance of CD4⁺ T cells in expanded TIL products (Fig. 1C) and their trend toward enrichment in TIL⁺ tumors, we performed CD4⁺ T cell–centric neighborhood analysis using our established pipeline. Windows of 20 nearest neighbors around each CD4⁺ T cell were clustered into 16 compositionally distinct T cell–centric neighborhoods (TCNs; Fig. 4G–I). TCN2, enriched for CD163⁺CD206⁺ macrophages, was nearly absent in TIL⁺ tumors, whereas TCN3, enriched for CD8⁺ T cells and CD68⁺ macrophages, trended toward higher abundance in TIL⁺ tumors, although these differences were not significant after two-sided Mann–Whitney U testing with BH-FDR correction (Fig. 4J, K; Supplementary Fig. 5C). The findings suggest that TIL⁺ and TIL⁻ tumors contain distinct immune compartments that may differentially support or suppress TIL expansion.

We next assessed whether these compositional differences were accompanied by altered spatial proximity using cell–cell interaction analysis, comparing observed cell-type distances with a null distribution generated from 1,000 random label permutations. Among interactions with log fold-change >0.1, CD4⁺ T cells showed group-specific coupling to macrophage subsets. In TIL⁺ tumors, CD4⁺ T cells were positioned closer to CD68⁺ macrophages, whereas in TIL⁻ tumors they were more closely associated with CD163⁺CD206⁺ macrophages (Fig. 4D, L). Thus, T cell positioning appears to reflect selective spatial coupling with functionally distinct macrophage populations.

Transcriptional analyses of myeloid compartments supported these spatial findings (Supplementary Fig. 5D–F). In TIL⁺ tumors, microglia-like TAMs were enriched for inflammatory and cytokine-responsive pathways, including TNFα/NF-κB, IL2/STAT5, and inflammatory response signatures. Monocytes showed similar cytokine-responsive and metabolically active programs, consistent with recruitment and activation. Bone marrow–derived TAMs in TIL⁺ tumors were enriched for inflammatory, complement, antigen-presentation, and proliferative pathways, whereas TIL⁻ tumors showed hypoxia-associated signaling, consistent with a metabolically constrained, immunoregulatory phenotype.

Our data indicate that macrophage abundance differs only modestly by TIL status, whereas macrophage state and spatial coupling to T cells diverge substantially. TIL⁺ tumors show T cell proximity to inflammatory and antigen-presenting macrophage populations, while TIL⁻ tumors show preferential T cell association with macrophages displaying hypoxia-associated and immunoregulatory programs. These findings support a model in which myeloid context and spatial organization, rather than abundance alone, shape the local immune microenvironment and influence TIL expansion potential.

## Discussion

We integrated ex vivo TIL expansion with single-cell, spatial transcriptomic, and multiplexed protein profiling to examine glioblastoma immune architecture and its relationship to recoverable, expandable T cell states. Our data implicate the organization and functional context of the TME as key determinants of TIL expansion.

Across scRNA-seq, Xenium, and CODEX, glioblastomas showed a conserved but heterogeneous cellular architecture comprising malignant, lymphoid, myeloid, and stromal compartments. These populations were consistently detected across modalities, and their relative abundances varied across samples without segregating by TIL expansion status. In contrast, spatial analyses showed that immune cells are not uniformly distributed but occupy structured microenvironments defined by vascular proximity, tumor regions, and myeloid context. This organization was reproducible across patients and platforms, indicating that glioblastoma contains a conserved spatial framework within which immune variation occurs^16^.

In this context, a central contribution of this study is the definition of this spatial and functional framework itself. Across all samples, we identify recurrent features of glioblastoma immune organization, including vascular-associated immune niches, structured lymphoid–myeloid interactions, and spatially constrained immune cell distributions. These features were observed across tumors irrespective of TIL expansion status, indicating that they likely reflect recurrent spatial programs of the glioblastoma microenvironment that can be variably leveraged, rather than niche properties confined to a distinct subset of tumors.

Within this shared architecture, we demonstrate that TIL generation from glioblastoma is feasible but variable. Expansion yielded products enriched for antigen-experienced T cells, predominantly of an effector memory phenotype, with oligoclonal TCR repertoires and preserved cytokine production following stimulation. These features are consistent with prior antigen exposure and suggest that at least a subset of expanded cells derive from tumor-resident clonotypes. However, variability in expansion efficiency, phenotype, and functional responsiveness was not explained by overall immune infiltration or the abundance of specific immune subsets in the tumor, indicating that bulk measures of immune presence are insufficient to predict TIL yield.

Instead, differences between TIL⁺ and TIL⁻ tumors emerged at the level of spatial relationships and intercellular connectivity within this common framework. TIL⁺ tumors were characterized by closer lymphoid–endothelial proximity, increased representation of vascular-associated immune niches, and enhanced endothelial-centered interaction networks inferred from CellChat. These findings point to a microenvironment in which endothelial structures act as organizational hubs for immune cells, potentially facilitating local interactions that support T cell survival, activation, or retention. Complementing this, lymphoid cells in TIL⁻ tumors were more frequently enriched within a necrosis-associated niche, suggesting that immune cells may be present in some non-expanding tumors but preferentially localized to regions less supportive of productive T cell persistence or recoverability.

The myeloid compartment further refines this organization. Although overall macrophage abundance differed only modestly, T cell–macrophage spatial coupling was distinct. In TIL⁺ tumors, CD4⁺ T cells were preferentially positioned near CD68⁺ macrophages, whereas in TIL⁻ tumors they were more closely associated with CD163⁺CD206⁺ macrophages. Transcriptional analyses supported this distinction, with TIL⁺ tumors enriched for inflammatory and cytokine-responsive programs across microglia-like TAMs, monocytes, and bone marrow–derived macrophages, while TIL⁻ tumors showed hypoxia-associated programs. These findings suggest that myeloid functional state and spatial relationship to T cells may influence whether T cells occupy environments permissive or restrictive for activation and expansion^17^.

Critically, these differences are not absolute. Considerable overlap exists between groups and canonical chemokine pathways such as CXCL12–CXCR4 show heterogeneous spatial expression without consistent enrichment in either condition. This argues against a single dominant signaling axis and instead supports a model in which multiple interacting features, including vascular proximity, myeloid state, regional tissue stress, and local cell–cell interactions, define the immune context.

Our findings shift interpretation of TIL biology in glioblastoma away from a binary classification of “inflamed” versus “non-inflamed” tumors. Many glioblastomas appear to contain T cells with the capacity for expansion, but this potential is variably realized depending on how these cells are positioned and conditioned within the TME. This is consistent with evidence across cancers showing that effective antitumor T cell immunity can depend on specialized perivascular immune niches, where vascular, dendritic, and lymphoid interactions coordinate local immune activation and restraint^18^. Accordingly, TIL⁺ and TIL⁻ tumors do not represent distinct biological categories, but rather different points along a continuum of immune organization, vascular niche support, regional exclusion into necrotic or hypoxic zones, and immune accessibility.

This interpretation is further supported by broader spatial studies in glioblastoma demonstrating that myeloid populations occupy reproducible anatomical compartments shaped by regional hypoxia, chemokine gradients, and homotypic and heterotypic cellular interactions^19,20^. Likewise, glioblastoma cells show recurrent clustering and dispersion across cohorts and platforms, with clustering linked to stabilized tumor cell identity states and dispersion to increased plasticity. Together, these findings suggest that spatial organization is a fundamental feature of glioblastoma biology with practical therapeutic implications. If TIL expansion potential is shaped by a shared but variably configured microenvironment, strategies may need to focus less on increasing immune cell abundance and more on remodeling local tissue architecture, including enhancing vascular–immune coupling, restoring supportive perivascular niches, limiting immune sequestration in necrotic or hypoxic regions, and reprogramming suppressive myeloid states toward inflammatory or antigen-presenting phenotypes.

Several limitations should be considered. Spatial and cellular heterogeneity is inherent to resected glioma specimens and may contribute to variability across samples. The ex vivo expansion protocol imposes selective pressures that may alter T cell composition relative to the *in situ* state, and expansion was not normalized to initial T cell input. The cohort size was limited, comparisons between TIL⁺ and TIL⁻ tumors were not powered for definitive statistical inference, and the analyses were correlative. Assessment of tumor antigen specificity was not performed, and not all assays were available for all samples, limiting cross-modal integration.

Nonetheless, our study provides a multimodal view of the glioblastoma immune microenvironment by integrating cellular composition, spatial organization, and functional state. Our data suggest that determinants of TIL expansion are not categorical, but are embedded within the architecture of a shared yet variably organized tumor ecosystem. T cell positioning relative to endothelial structures, together with the state and localization of myeloid populations, may influence whether antitumor immunity is supported or restrained. T cells within organized perivascular niches may receive trafficking, survival, and activation cues, whereas those confined to necrotic, hypoxic, or suppressive regions may remain dysfunctional despite being present. Similarly, inflammatory myeloid populations near vascular interfaces may support immune surveillance, while suppressive macrophage states may impede it.

More broadly, emerging studies indicate that spatial transcriptomic approaches can improve biologic and diagnostic resolution in brain tumors^21^. Our findings support a model in which the TME itself is a therapeutic target. Strategies that enhance vascular–immune interactions, relieve hypoxia-driven suppression, reduce immune exclusion, or redirect myeloid cells toward inflammatory states may improve the effectiveness of T cell–based therapies in glioblastoma.

## Acknowledgements

General

The authors are grateful to all patients who consented to donate biospecimens for this study. We thank the team at the Duke Brain Tumor Biorepository for their assistance and co-ordination in tissue management and slide preparation. We specially thank Drs. Peter Fecci, Anoop Patel and Allan Friedman, the neurosurgeons who provided the surgical specimens. We would also like to acknowledge the assistance of the Molecular Genomics Core at the Duke Molecular Physiology Institute, Duke University School of Medicine, for the generation of data for the manuscript.

## Funding

NIH Duke SPORE (P50 CA190991-08) DRP grant (MK and KMH)

Knox Martin Foundation (MK)

Barzun and Siebert Funds (MK)

Viral Oncology T32 (T32CA009111) (KMH)

## Institutional Review Board (IRB) / ethics approval

All human samples were collected on Duke IRB protocol number Pro00106078.

## Author contributions

Conceptualization: KMH, PN, JWH, MK

Methodology: KMH, KZ, AMC, EO, JWH, MK

TIL expansion: PN, YZ, BS

scRNAseq library preparation: CR, KMH

Xenium Slide preparation: Duke Brain Tumor Biorepository

Xenium raw data generation: The Molecular Genomics Core at the Duke Molecular Physiology Institute, Duke University

Custom Xenium Panel Design: KMH, AMC, JCG, MCB, KA, MK

scRNA analysis: EO

spatial RNA Xenium data analyses: AMC, KZ, MK

Bulk RNA/ TCRseq: KZ, MK

Codex: JWH, KVB

Flow cytometry: KMH

H&E slides and images review: RM

Writing – original draft: KMH, EO, AMC, KZ, JWH, MK

Writing – review & editing: all authors

Supervision: MK

## Online methods

### Sample preparation

The glioma samples were collected as per Duke Institutional Review Board approved protocol “Collection of Solid Tumor Specimens for Pre-clinical Development and Manufacturing of a Clinical Tumor Infiltrating Lymphocyte (TIL) Cell-therapy Product” Pro00106078 for the collection of the preclinical development samples. Patients were consented for biobanking of their tumor and blood specimens, sample acquisition and data collection.

### TIL expansion

Autologous TILs were generated in the Process Development Laboratory at the Marcus Center for Cellular Cures, Duke University. The preREP/REP protocol, originally described by Steven A. Rosenberg and colleagues, was adapted by the Marcus Center for Cellular Cures (MC3) team at Duke University using lung tumor–derived tumor-infiltrating lymphocytes (TILs) to streamline manufacturing and reduce cost for clinical-scale manufacturing^10^. Here, preREP refers to the pre-rapid expansion phase, and REP refers to the rapid expansion phase. Feeder cells were replaced with CD3/CD28 activation beads, eliminating the need for extensive feeder validation and reducing labor. Although feeder-based expansion yielded higher cell numbers, feeder-free conditions produced TILs with improved cytotoxicity, consistent with prior reports showing preferential outgrowth of non–tumor-specific bystander clones during REP^22^. Despite lower expansion rates, feeder-free REP generated clinically sufficient yields (mean 1.4×10¹¹ cells per 0.5 g tumor; n=11).

Tumor fragments distinct from those used for multimodal profiling were processed to preserve native tumor characteristics. This approach was intentionally implemented to preserve the integrity of the original tumor specimen for downstream immune and spatial characterization, ensuring that *ex vivo* culture did not alter the molecular or cellular features assessed across profiling platforms. Fresh specimens were weighed, minced into 2–3 mm³ fragments, and cultured in G-Rex flasks (5–24 mg/cm²) in preREP media (PrimeXV, 5% human platelet lysate, 4,000 IU/mL IL-2, penicillin/streptomycin) for 18–28 days. Cells were harvested, counted, and cryopreserved (CryoStor CS10). For REP, TILs were thawed and cultured at 1×10⁶ cells/mL/cm² in REP media (PrimeXV, 3% human platelet lysate, 1,000 IU/mL IL-2) with CD3/CD28 activation (TransAct). Activation was halted after 3 days by dilution, with media exchanges guided by lactate levels (15 mM threshold). Cells were harvested at day 14 and cryopreserved.

Other reagents and materials included PrimeXV XSFM T-cell expansion media (Fujifilm, Santa Ana, CA), G-Rex flasks (Wilson-Wolf, St. Paul, MN), human platelet lysate (Compass Biomedical, Hopkinton, MA), penicillin/streptomycin (Gibco, Thermo Fisher Scientific, Waltham, MA), IL-2 (Bio-Techne, Minneapolis, MN), CryoStor CS10 (BioLife Solutions, Bothell, WA), CD3/CD28 nanoparticles (TransAct, Miltenyi Biotec, Bergisch Gladbach, Germany), and lactate measurement system (Lactate Plus meter and strips, Nova Biomedical, Waltham, MA).

### Flow cytometry

Cryopreserved cells were rapidly thawed in a 37°C water bath and resuspended in RPMI media with 2% fetal bovine serum. Cells were then pelleted at 300xg for 5 minutes at 4C and resuspended to count by hemocytometer. Cells for immune phenotyping one million cells per well were loaded into a round bottom 96 well plate and stained immediately post thaw. For TIL stimulation assays cells were rested overnight in an incubator at 37°C and 5% CO2. Post rest cells were counted and plated in tissue culture treated polypropylene 96 well plates at one million cells per well. TILs were stimulated with either 25uL CD3/CD28 stimulation (ImmunoCult™ Human CD3/CD28 T Cell Activator Catalog#10971, StemCell) or 1x concentration of PMA/ion (eBioscience™ Cell Stimulation Cocktail (500X), Catalog# 00-4970-93, ThermoScientific) per million cells in 200uL media for 6 hours. Matched unstimulated TILs were cultured in media for 6 hours at 37°C and 5% CO2. Degranulation marker antibody, CD107a-PE, was added during cell stimulation along with brefeldin (BD GolgiPlug) and monensin (BD GolgiStop). Following the 6 hour stimulation, the cells were washed to remove media and stimulation cocktails. Cells were first stained with LiveDead Blue Fixable viability dye (Invitrogen), then blocked with buffer containing anti-CD16/32 (Invitrogen) and mono-block (BioLegend). A master mix containing all surface antibodies was prepared in brilliant violet stain buffer (Invitrogen) and added on top of blocking buffer for 30 mins. Cells were then fixed and permeabilized using the eBioscience™ Foxp3 / Transcription Factor kit to allow for intracellular staining of cytokines and intracellular markers. Spectral data was collected on the Cytek Aurora 5L and analyzed by FlowJo v10.

Details of flow cytometry antibodies used are in the supplementary material (Supplementary Table 2).

### Sample preparation for multimodality analyses

Formalin-fixed paraffin-embedded (FFPE) tissues were processed for single-cell and xenium profiling, followed by bulk TCRseq and CODEX multiplexed imaging.

#### Formalin-Fixed Paraffin-Embedded (FFPE)

5mm FFPE tissue sections are placed on a 10x Genomics Xenium slide. Tissue slides are deparaffinized and decrosslinked. Next, tissue is processed through Hybridization, Ligation & Amplification protocols. Prepared tissue slides are then loaded for imaging on the Xenium Analyzer for *in situ* analysis. Cyclical rounds of fluorescent probe hybridization, imaging, and removal generate optical patterns specific for each barcode, which are converted into a gene identity. Identified transcripts are then visualized using Xenium Explorer software. H&E staining was performed on the slides following Xenium analysis to allow for histopathological assessment.

#### FFPE with custom panel and multimodal segmentation

5um FFPE tissue sections are placed on a 10x Genomics Xenium slide and baked at 42C for 3hours. Once dried, tissue slides are stored in a desiccator to await processing. Xenium slides are then incubated at 60C for 2 hours and then deparaffinized and decrosslinked. Briefly, sections are deparaffinized via xylene immersion to solubilize the paraffin, ethanol to remove the solubilized paraffin and slowly hydrate the tissue, and then immersed in water to remove residual ethanol. Once hydrated, tissues slides are decrosslinked. Tissue slides are inserted into 10x Genomics Xenium slide cassettes. Decrosslinking buffer is then added to the well created by the cassette and incubated. After incubation, the buffer is removed, and the sample is washed. Tissue is then processed through Hybridization, Ligation & Amplification protocols. Briefly, a custom panel containing probes for 480 custom gene panel *(The gene list is available as a supplementary file)* enriched in genes relevant to glioma, neuronal and immune cells, is added to the tissue. The 480-gene panel was used to enable mapping of gene expression in the tissue architecture. Differential expression analyses highlighted immune-tumor dynamics in pro-inflammatory and immunosuppressive pathways. Each circularizable DNA probe contains two regions that hybridize to target RNA and a third region that encodes a gene-specific barcode. The two ends of the probes bind the target RNA and are ligated to generate a circular DNA probe. Following ligation, the circularized probe is amplified, producing multiple copies of the gene-specific barcode for each target. Tissue slides are then stained using the 10x Genomics Multimodal Cell Segmentation Kit. During Multimodal staining, antibodies used for cell segmentation bind their antigens in an overnight incubation, followed by post-incubation washes to remove excess antibodies. The cell segmentation stain is then enhanced by 10x Genomics proprietary reagents. The Multimodal Cell Segmentation kit stains for cell nuclei, membranes, and cell interior that are inputs for the 10x Genomics automated morphology-based cell segmentation analysis pipeline. Prepared tissue slides are then loaded for imaging on the Xenium Analyzer for *in situ* analysis. Fluorescently-labeled oligos bind to the amplified DNA probes. Cyclical rounds of fluorescent probe hybridization, imaging, and removal generate optical patterns specific for each barcode, which are converted into a gene identity. Identified transcripts are then visualized using Xenium Explorer software.

### Chromium single cell gene expression flex (scFFPE-seq)

For each sample, 2x 25 μm FFPE curls were dissociated in accordance with 10x Genomics “Isolation of Cells from FFPE Tissue Sections for Chromium Fixed RNA Profiling protocol (User guide CG000632, Rev. B). Following cell isolation, samples underwent probe hybridization with one probe barcode per sample from the 10x Genomics Single Cell Fixed RNA Human Transcriptome Probe Kit. Approximately 151,050 cells per sample were pooled, washed, filtered, and loaded into 10x Genomics’ Next Gem Chip Q targeting a recovery of 8,000 cells per probe barcode. Sequencing libraries were generated following 10x Genomics’ User Guide CG000527 for Fixed RNA Profiling (Rev. E). The tumor samples from eight patients were used to generate two libraries following 10x Genomics’ User Guide CG000527 for Fixed RNA Profiling (Rev. E). These libraries were sequenced on an Illumina NovaSeq platform, resulting in four FASTQ files per library. Individual samples in each library were demultiplexed by their incorporated barcodes using CellRanger’s (v7.0.1) multi pipeline and the GRCh38-2020-A reference (10x Genomics). Following demultiplexing, sequencing data from each sample was obtained, enabling data analysis across both libraries. CellRanger’s aggr pipeline was utilized to aggregate data from the same patients, treating data from different libraries as independent replicates to evaluate any biological variability and to provide more substantial cell counts.

### Single-cell RNA sequencing (scRNA-seq)

CellRanger’s pipelines corrected for empty droplets, cells with few or no transcripts, an important variable in quality control filtering. Cells were further filtered to ensure high quality reads, which including removing ambient RNA and potential doublets. DoubletFinder was used to estimate and remove doublets from the samples, using 10x Genomics’ Fixed RNA Profiling protocol’s predicted doublet rate of 0.8% per 1,000 cells obtained as a parameter. Additionally, cells with greater than 10,000 detected transcripts, greater than 7,000 detected genes, a mitochondrial gene proportion exceeding 20%, a cell complexity below 80% (log10Gene/log10UMI), and a hemoglobin gene proportion above 2% were excluded before downstream analysis. The hemoglobin gene threshold of 2% was stringent since only a few cells across the eight samples exceeded this level, reflecting the solid tissue origin of the samples. Genes present in fewer than three cells per sample were also removed to eliminate low-abundance genes and correct potential misreads from the reference transcript. Although a threshold of 30 cells is commonly used, this was adjusted for the low cell count of these samples, ranging from 539 to 1429 cells. Ribosomal gene proportions, another common parameter, were not applicable because CellRanger’s reference transcriptome excludes ribosomal reads. Violin plots confirmed that filtering did not result in data loss, but this filtering ensures that downstream analysis will not be influenced by technical variations or losses. In total, 6,567 cells were included in the downstream analysis, with 3,897 cells in the TIL^+^ group and 2,670 cells in the TIL⁻ group.

After normalization and scaling with Seurat, the data was batch-corrected and integrated using Seurat’s integration pipeline. The IntegrateLayers function applies canonical correlation analysis (CCA), a method akin to principal component analysis, to identify shared sources of variation between sample groups (TIL^+^ vs. TIL⁻). Jackstraw plots and clustree were used to determine the appropriate number of principal components and value of resolution, respectively, before running this analysis. This approach removes non-biological variation, accounts for confounding factors, and enables comparative analysis between groups. To ensure reproducibility, a random seed was set for Seurat’s RunPCA, FindClusters, and RunUMAP.

Post-integration, clusters identified using Seurat’s integration method were automatically annotated using the SingleR package, referencing a single-cell RNA sequencing glioblastoma dataset from CELLxGENE. SingleR^23^, a correlation-based method, matches the full transcriptome of query cells to reference cells rather than relying solely on marker gene expression. Annotation accuracy was assessed by identifying differentially expressed genes (DEGs) for each cluster using Seurat’s FindMarkers function, comparing each cluster against the remaining cells. Tumor subtypes were validated based on prior literature^24^, with AC-like clusters expressing GFAP, BCAN, SPARC, and DBI; MES-like clusters expressing HILPDA, CA9, ADM, and VIM; NPC-like clusters expressing DCX, DLX5, SOX4, DBN1, and RND3; and OPC-like clusters expressing NEU4, PTPRZ1, LIMA, VCAN, and OLIG1. Non-malignant clusters were further validated using marker genes from CellMarker2.0^25^ and published literature^26^: astrocytes (GFAP, CLU, AQP4, S100B), CD4/CD8 T cells (CD3E, CD3D, CD3G, CD4, CD8A/B, GMZA, SKAP1), endothelial cells (VWF, CLDN5, CLEC14A, CDH5), mural cells (CD248, PDGFRB, RGS5, FOXF2), neurons (SYT1, SNAP25, GABRA1, NRCAM), oligodendrocytes (MOG, MBP, OLIG1, OLIG2, CLDN1), OPCs (SOX10, PDGFRA, APOD, OLIG1, OLIG2), TAM-BDM (CD163, CD14, TGFBI, ITGA4, MARCO), TAM-MG (TMIGD3, APOC2, SCIN, P2RY12, NAV3), and monocytes (LYZ, AREG, CD163, CD68, MXD1).

Differential gene expression (DGE) analysis was conducted between TIL^+^ and TIL⁻ groups using EdgeR, which is a Bioconductor software package for examining differential expression of replicated count data^27^. These methods were also applied to identify differentially expressed genes among cell types within each group. Cell composition analysis was performed between the two groups to identify the proportion of each cell type in the TIL^+^ vs TIL⁻ samples. The Mann-Whitney U test was used to determine whether the differences in cell composition between the sample groups were significant. Violin plots were also made to illustrate sample variability in cell compositions.

### Xenium in situ spatial transcriptomics

Xenium sample results were read using the *ReadXenium* function, and the centroids, segmentations, and coordinates of each sample were added to a Seurat object. Transcripts with locations that did not overlap nuclei coordinates or with quality value (qv) scores ≤ 20 were filtered out. The probes for FLT3LG (FLT3 ligand) and CCL5 were removed from the data due to high sensitivity and low specificity as confirmed by 10X Genomics. Then, cells were filtered to contain more than 5 features and more than 25 total counts. This filtered out a median of 12.39% of cells across the 11 samples. The samples were merged into one spatial experiment object, *computeLibraryFactors* was run, and then it was converted to a Seurat object for analysis. The standard Seurat pipeline was run on the merged object (*NormalizeData, FindVariableFeatures*, *ScaleData*, *RunPCA*, *FindNeighbors*, *FindClusters* (with resolution = 0.08 and 0.2), *RunUMAP*).

The “EGFR-exon1-exon2”, “EGFR-exon1-exon8”, “HERV-K-Gag”, and “HERV-K-Env” features were removed to prevent clustering based on EGFRv3 type tumors, as the goal was to split tumor cells by subtypes, and to avoid clustering by virus expression. Additionally, the “PDCD1”, “NCSTN”, “SCD”, “SPOCD1”, “GFAP”, “ACTA1”, “PVR”, “PPARG”, “CNTN2”, “KLF2”, “TENM1”, and “CCL19” probes were removed from clustering due to high sensitivity and low specificity. Batch effect was not visualized in a UMAP by sample run. *Clustree* was used to select a clustering resolution of 0.2, and initial annotations were made based on marker genes, DGEs, and clustering. From this, immune cells were subset from the object and clustered using the same pipeline as the whole object. This generated six clusters, which were annotated based on marker gene expression. One sample showed low gene expression variability with largely nonspecific signal. Pathologic review confirmed that the tissue had been affected by electrocautery, and the sample was therefore excluded from the Xenium analysis.

Tumor cells were analyzed using GeneNMF^28^, which extracts common characteristics in cancer cells across different samples. It uses non-negative matrix factorization (NMF) to break down each sample’s expression matrix into a low number of dimensions, and then identifies meta-programs (MPs) that are expressed across samples. These MPs are characterized as consensus gene sets. The tumor cells were split by sample, then *multiNMF* was run with k=5:7 and min.exp = 0.05. Next, *getMetaPrograms*was run, and 10 MPs were found. Within these, 2 had low silhouette and mean similarity so they were dropped. Module scores were computed for each cell using *AddModuleScore_UCell,* and cells were assigned the MP with the highest score. Heatmaps of sampled cells vs MPs were plotted to evaluate performance, and marker genes of each MP were analyzed to annotate cell types. Subtypes were annotated based on marker gene expression and were distributed across samples, demonstrating distinguished cell types.

Xenium niches were computed spatially using *RunBanksy*^13^ with k = 30 neighbors and lambda = 0.8. Cell type niches were computed using Seurat’s *BuildNicheAssay* with k = 30 neighbors and k = 6 niches. Niche compositions were compared based on the cell types of neighboring cells.

Endothelial contour analysis was performed using Scider^29^ following the official guidelines (V 1.7.0). Tissue slides were segmented into 10μm hexagonal spatial bins, and cell-type densities were estimated by Kernel density estimation (KDE). Density correlations between cell types were quantified with Pearson correlation, restricted to the top 50% high-density spatial bins. Epithelial contour levels were defined by the boundaries of regions at different cell-density levels, with each level containing comparable target cell counts.

Ambient gene expression was estimated using *RunBanksy* based on the nearest 15 cells using a λ value of 0.8. Differentially expressed genes were defined by a fold change > 2 and a p-value < 0.05. Gene set enrichment analysis (GSEA) was performed with fgsea (V1.32.4)^14^. Spatial cell interaction was inferred using CellChat v2 (V2.1.0)^12^, and interaction events were quantified within a 10μm neighborhood. Visualization was generated using the default parameters.

### CODEX multiplexed imaging and data

#### Array creation

Imaging data was collected from multiple brain samples from multiple donors on one slide. We included up to six tissues that were cut onto the same slide with a 4 μm section thickness and mounted onto Superfrost PLUS slides for further processing.

#### Antibody conjugation

Each antibody was conjugated to a unique oligonucleotide barcode using our previously established protocol^6^, after which the tissues were stained with the antibody–oligonucleotide conjugates and we validated that the staining patterns matched the expected patterns already established for immunohistochemistry within positive control tissues of the intestine or tonsil. Otherwise, expected co-expression of markers in the case of staining of suspension cells was considered for validating separate cell type and phenotype antibodies. First, antibody–oligonucleotide conjugates were tested in low-plex fluorescence assays and the signal-to-noise ratio was also evaluated at this step, then they were tested all together in a single CODEX multicycle.

#### CODEX staining and imaging

CODEX multiplexed imaging was executed according to the CODEX staining and imaging protocol.^6^ Briefly, the tissue arrays were stained with the validated panels of CODEX antibodies and imaged using Akoya PhenoCycler Fusion 2.2.0, involving cyclic stripping, annealing, and imaging of fluorescently labeled oligonucleotides complementary to the oligonucleotide conjugated to the antibody. Each slide underwent CODEX multiplexed imaging using Akoya PhenoCycler Fusion machine; metadata from each CODEX run can be found in the supplementary files (Supplementary Table 3).

#### CODEX data processing

Raw imaging data were processed using the Akoya PhenoCycler Fusion 2.2.0 software for image stitching, drift compensation, deconvolution, and cycle concatenation. Processed mages were visualized in napari (https://github.com/napari/napari) and evaluated for specific signal. Any markers that produced an untenable pattern or a low signal-to-noise ratio were excluded from the ensuing analysis. To obtain quantitative single cell information, processed images were then segmented using Mesmer, a deep-learning based tissue segmentation algorithm, via SPACEc Python package, which is available for download from https://github.com/yuqiyuqitan/SPACEc.

#### Cell type analysis

A total of 2,403,156 cells were identified across 11 samples and classified into 21 cell types and states based on marker expression. Cell type identification was done following the strategies we have developed (citation). Briefly, nucleated cells were selected by gating DAPI-positive cells, followed by z-normalization of protein markers used for clustering (some phenotypic markers were not used in the unsupervised clustering). The data were overclustered with Leiden-based clustering with the scanpy Python package. Clusters were assigned a cell type based on average cluster protein expression and location within image. Impure clusters were split or reclustered following mapping back to original fluorescent images.

#### Cellular neighborhood analysis

Neighborhood analysis was performed as described previously^30,31^. In brief, this analysis involved (1) taking windows of 20 cells across the entire cell type map of a tissue with each cell as the center of a window; (2) calculating the number of each cell type within this window; (3) clustering these vectors using k-means clustering; and (4) assigning overall structure on the basis of the average composition of the cluster. We initially generated 25 clusters; these clusters were then assessed for both cellular composition and spatial localization. Clusters that were spatially co-localized and exhibited similar cell type compositions were subsequently merged, resulting in 13 distinct cellular neighborhoods.

#### CD4^+^ T cell-centric neighborhood analysis

CD4^+^ T cell-centric neighborhood analysis was performed as described previously^11^. For CD4^+^ T cell-centric neighborhoods, we used the cell neighbor composition vectors by taking the 20-nearest neighbors surrounding the CD4^+^ T cells. We then clustered cell neighborhoods with k-means clustering. We initially generated 25 clusters; these clusters were then assessed for both cellular composition and spatial localization. Clusters that were spatially co-localized and exhibited similar cell type compositions were subsequently merged, resulting in 16 distinct CD4^+^ T cell-centric neighborhoods.

#### Cell interaction analysis

Cell interaction analysis was carried out as described previously^32^. Briefly, the Delaunay triangulation of cells were identified within each field of view using the x, y position. Cell to cell interactions within 128 μm from one another were identified and baseline distributions were established by performing the calculation for each iteration the cell label was randomly assigned existing x, y positions and compared to observed distances with a Wilcoxon Test. We used 1,000 permutations for the distribution. The fold enrichment of distances between the observed data over the mean distances from the permutation test. The log fold of the distances for each cell type interaction where p values less than 0.05 were plotted for each group.

### TCR repertoire analyses

T cell receptor sequencing (TCRseq) was performed using iRepertoire’s RepSeq+ technology to assess the diversity and clonality of immune repertoires. The diversity and clonality of tumor-infiltrating lymphocyte (TIL) populations were evaluated, with a specific focus on clonal expansions, particularly among CD8^+^ T cells in TIL^+^ tumors, to quantify tumor-reactive TCR frequencies.

#### TCRseq

RNA was extracted from TIL samples using the PAXgene RNA Extraction Kit (PreAnalytix), and total RNA were extracted from FFPE samples using the Covaris truXTRAC FFPE total NA Plus Kit, and SPRIselect beads were used for RNA concentration to maximize input for amplification. 1600 ng of RNA per sample, along with replicates, was utilized for RepSeq+ TCR alpha, beta, delta, and gamma chain amplification. The RepSeq+ protocol facilitated simultaneous amplification of all four TCR chains under uniform conditions using primer pairs specific to V-D-J combinations, incorporating unique molecular identifiers (UMIs) during reverse transcription to minimize sequencing errors and PCR duplicates. Sequencing was performed on the Illumina NextSeq 1000 platform using a P1 600-cycle kit.

Raw sequencing data were processed using the iRmap pipeline, in which sequence reads were de-multiplexed based on Illumina dual indices and barcode sequences, merged, and mapped to germline V, D, J, and C reference sequences using the IMGT reference library. CDR3 regions were then identified, extracted, and translated into amino acid sequences. The final dataset was refined by collapsing identical CDR3 and UMI combinations to correct for sequencing errors. TCR repertoire diversity and clonality were evaluated by D50, Diversity index, and Shannon entropy. The D50 is the percentage of dominant and unique T cell clones that account for the cumulative 50% of the total CDR3s counted. The diversity index is 100 minus the area under the curve between the percentage of total reads and the percentage of unique CDR3s.

### Statistical analysis

Statistical analyses were performed using modality-specific R and Python workflows described above. Unless otherwise stated, comparisons between TIL⁺ and TIL⁻ tumors were performed using two-sided non-parametric tests, with the patient/sample treated as the primary unit of analysis to avoid inflation from single-cell measurements. Paired comparisons of TIL yield between pre-REP and REP were performed on log10-transformed cell counts using a paired two-sided t-test, given that cell counts spanned several orders of magnitude. Differences in cell-type proportions, neighborhood abundance, lymphoid cell-state scores, endothelial proximity metrics, and CD4⁺ T cell-centric neighborhood features were assessed using Wilcoxon rank-sum/Mann–Whitney U tests, as appropriate. Multiple hypothesis testing was controlled using the Benjamini–Hochberg false discovery rate where indicated, particularly for neighborhood and high-dimensional feature comparisons. Differential gene expression between groups and within annotated cell populations was performed using edgeR, and pathway-level enrichment was assessed using fgsea-based gene set enrichment analysis. CODEX cell–cell interaction analyses compared observed spatial distances with null distributions generated from 1,000 random label permutations; interactions with nominal P < 0.05 and log fold-change > 0.1 were visualized. TCR repertoire metrics, including D50, diversity index, and Shannon entropy, were summarized descriptively. Given the limited cohort size and inter-patient heterogeneity, statistical analyses were considered exploratory and interpreted in the context of effect size, consistency across modalities, and biological plausibility rather than statistical significance alone.

## Data availability

Raw and processed data available on request.

## Code availability

TCRseq data, processed scRNA data and Xenium expression matrix with coordinates are available on GitHub at https://github.com/khasrawlab/TILproject

All demographic, flow cytometry data are available in the manuscript. Metadata and the custom gene panel is available as supplementary files.

Codex and Xenium images are available at FigShare

## Supplementary Figures

**Supplementary Figure 1.**
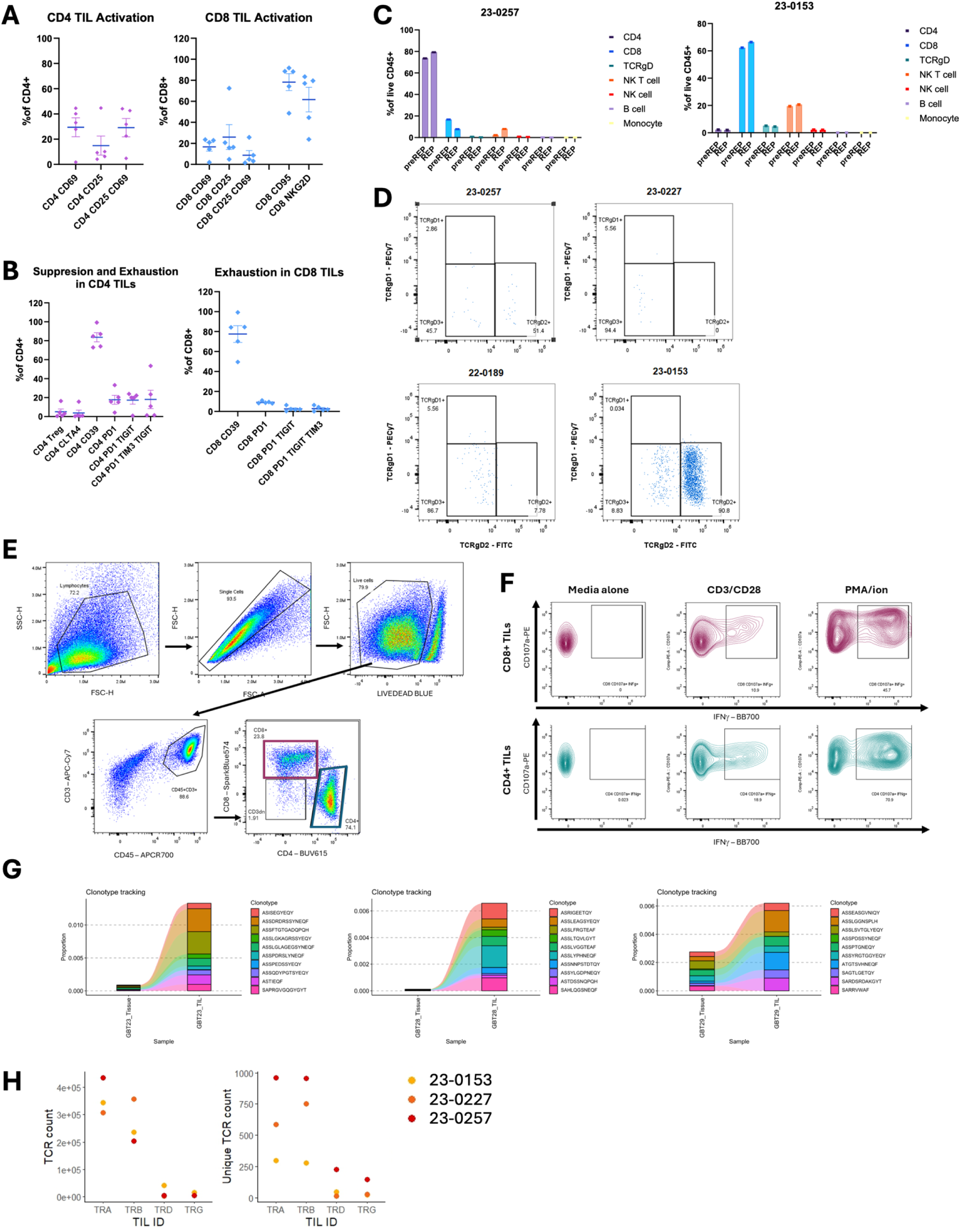
Immunophenotypic composition and TCR features of expanded TIL products. A, Activation markers. Activation markers were detectable in both CD4⁺ and CD8⁺ TILs, with inter-patient variability. B, Suppression and exhaustion markers. Markers associated with suppression or exhaustion were detectable at low levels in both CD4⁺ and CD8⁺ TILs, with variable expression across patients. C, Cell-type composition before and after expansion. Immune cell composition in representative samples (23-0257 and 23-0153) demonstrates heterogeneity in CD4⁺ versus CD8⁺ dominance and persistence of minor immune subsets between pre-REP and REP phases. D, γδ T-cell characterization. Representative flow cytometry plots showing variable TCRγδ subset distributions across selected samples. E, Intracellular cytokine gating strategy. Representative flow cytometry plots of cells cultured in media alone, stimulated with CD3/CD28, or stimulated with PMA–ionomycin. F, TCR repertoire tracking between tumor and expanded TILs. Frequencies of the ten most abundant shared TCRβ clonotypes in matched tumor tissue and expanded TIL products, demonstrating selective *in vitro* expansion of specific clones. G, Clonotype tracking analysis of T-cell receptor (TCR) repertoires between paired bulk tumor and REP TILs. H, Dot plots comparing TIL expansion across key clonality metrics (TCR count and unique TCR count).

**Supplementary Figure 2.**
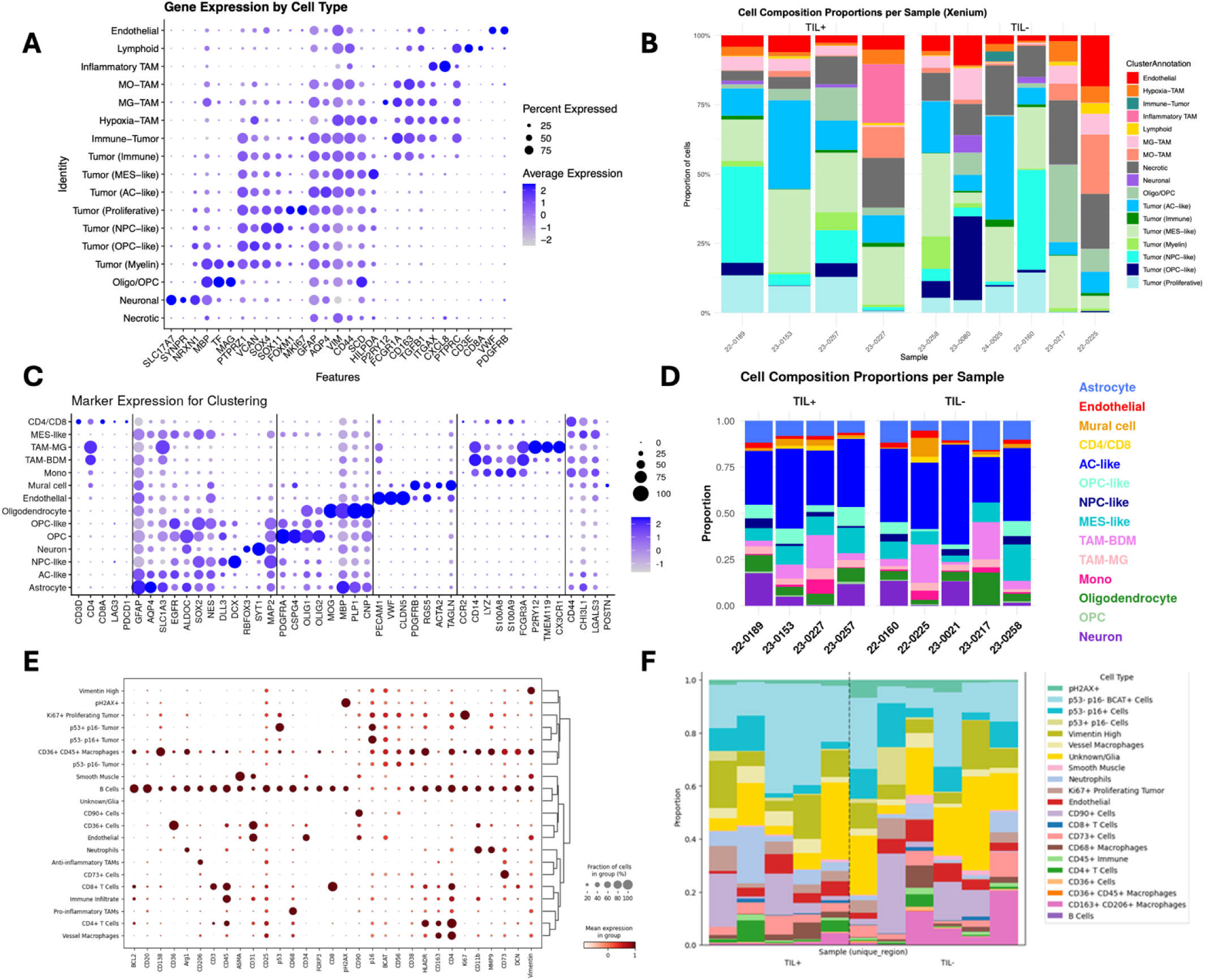
Cell type annotation and composition across Xenium, scRNA-seq and CODEX datasets. A, Xenium marker gene expression by cell type. Dot plot showing expression of canonical marker genes across annotated cell populations in the Xenium dataset. B, Xenium cell-type composition by sample. Stacked bar plots showing annotated cell-type proportions across individual Xenium samples. C, scRNA-seq marker gene expression by cell type. Dot plot showing marker gene expression across annotated scRNA-seq clusters. D, scRNA-seq cell-type composition by sample. Stacked bar plots summarizing cell-type proportions across samples stratified by TIL status. E, CODEX marker expression heatmap. Heatmap of protein marker expression across CODEX-defined cell populations with hierarchical clustering. F, CODEX cell-type composition by sample. Stacked bar plots showing proportions of CODEX-defined populations across samples.

**Supplementary Figure 3.**
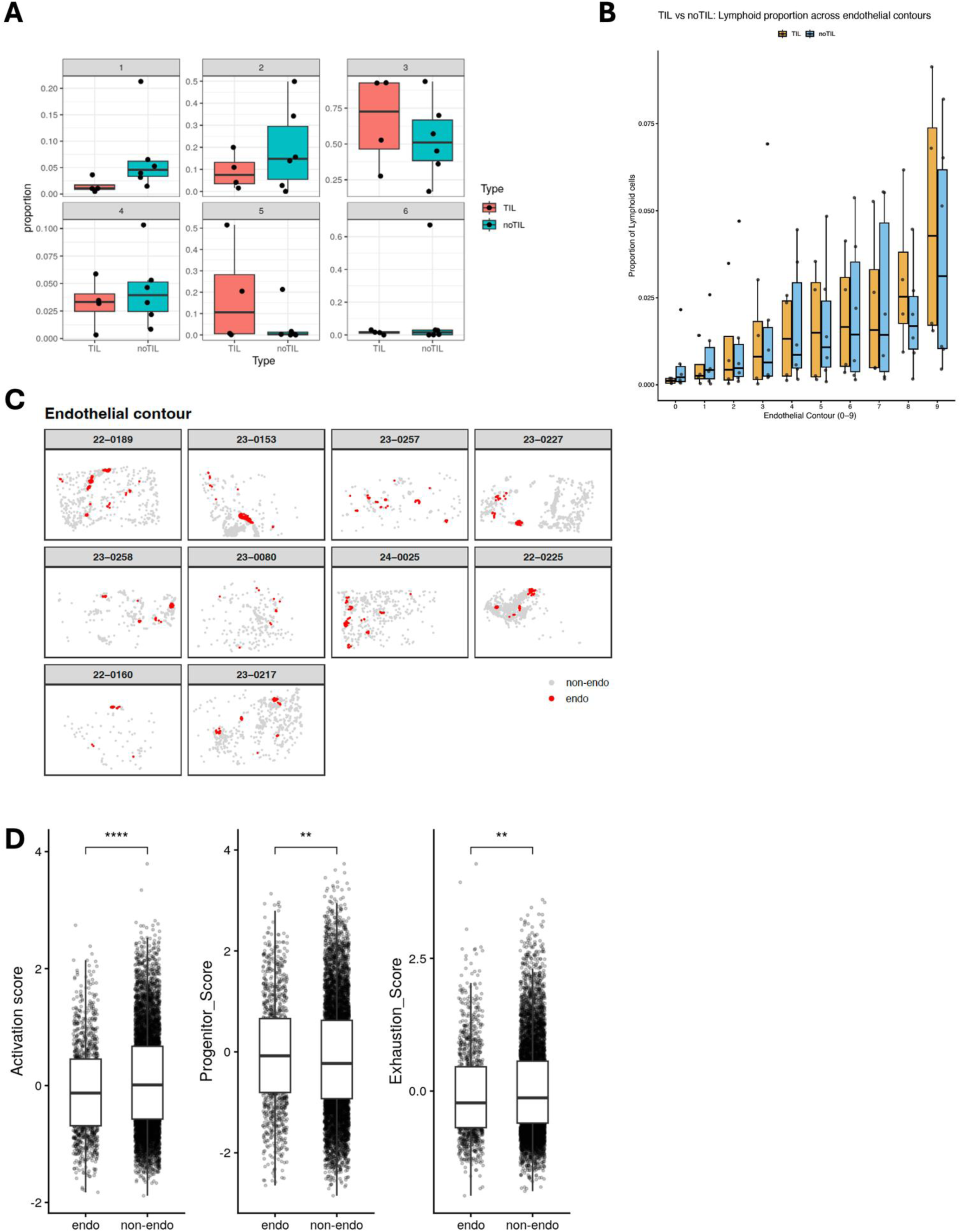
Spatial distribution of lymphoid cells and CXCL12–CXCR4 signaling. A, Lymphoid cell proportion across spatial niches. Boxplots comparing the proportion of lymphoid cells across defined niches between TIL⁺ and TIL⁻ tumors. B, Lymphoid cell proportion across contour levels. Distribution of lymphoid cell proportions across increasing contour levels, showing changes in density relative to local tissue structure. C, Spatial localization of lymphoid cells in high contour regions. Representative spatial maps highlighting lymphoid cells in high contour regions. D, Boxplots of lymphoid cell state scores derived from gene expression signatures, including activation, progenitor, and exhaustion-associated programs, were compared between lymphoid cells in or not in perivascular neighbors.

**Supplementary Figure 4.**
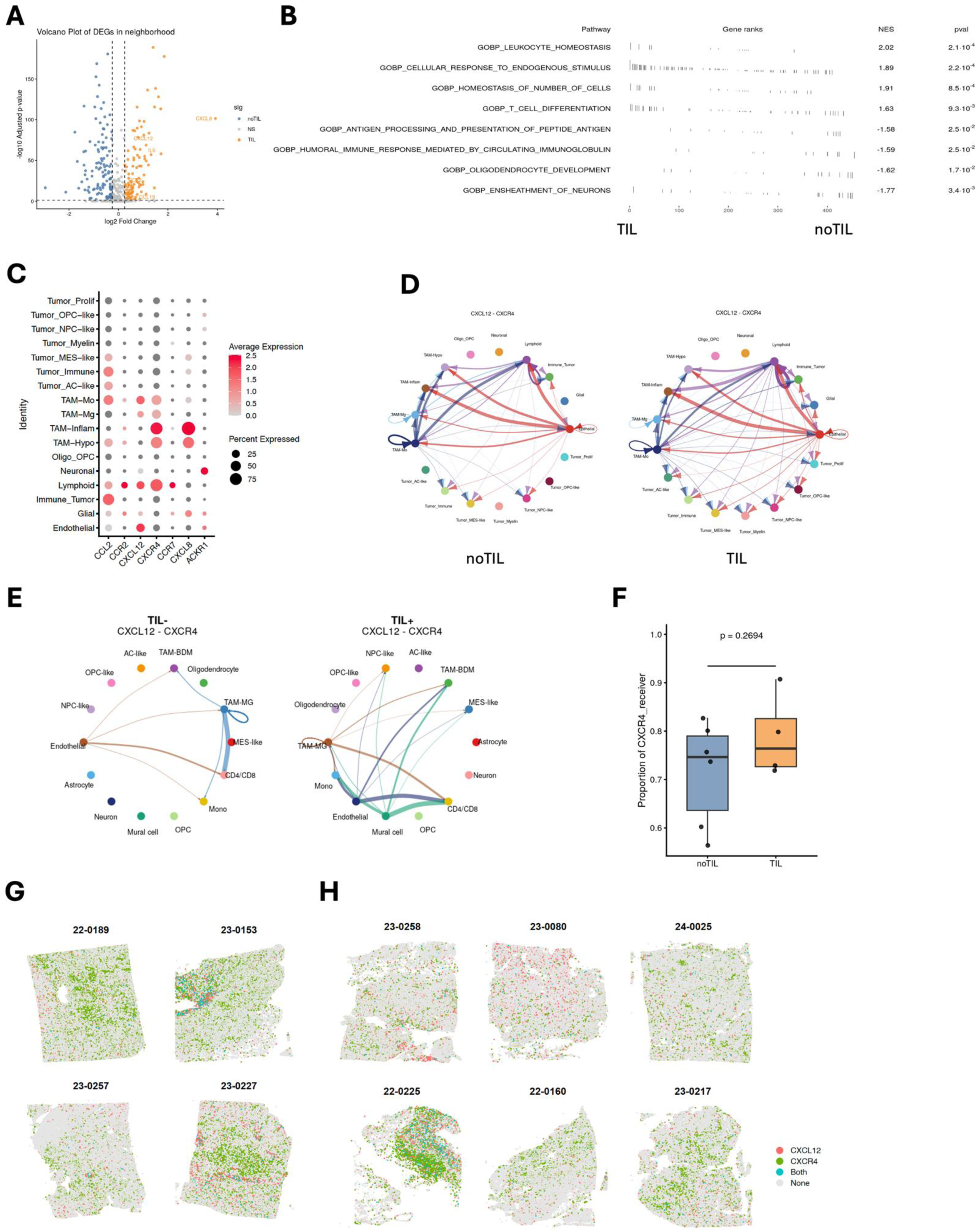
CXCL12–CXCR4 signaling shapes lymphoid cell states in TIL samples. A, Differentially expressed genes in lymphoid neighborhoods. Differentially expressed genes surrounding lymphoid cells in TIL and noTIL tumors, including multiple chemokines and cytokines enriched in TIL samples. B, Gene set enrichment analysis. Enrichment of immune-related biological processes in lymphoid neighborhoods from TIL samples. C, Ligand–receptor expression patterns. Dot plot showing expression of ligand–receptor pairs across cell clusters. D, Spatial cell–cell interaction analysis. Stronger CXCL12–CXCR4 interactions were inferred in TIL samples. E, Cell–cell interaction analysis in scRNA-seq dataset. Stronger CXCL12–CXCR4 interactions were inferred in TIL samples. F, CXCR4 receptor activity. Higher proportion of active CXCR4 receptors in lymphoid cells from TIL samples. G–H, Spatial distribution of CXCL12 and CXCR4 expression. Spatial maps showing heterogeneous CXCL12 and CXCR4 expression across TIL⁺ and TIL⁻ tumors.

**Supplementary Figure 5.**
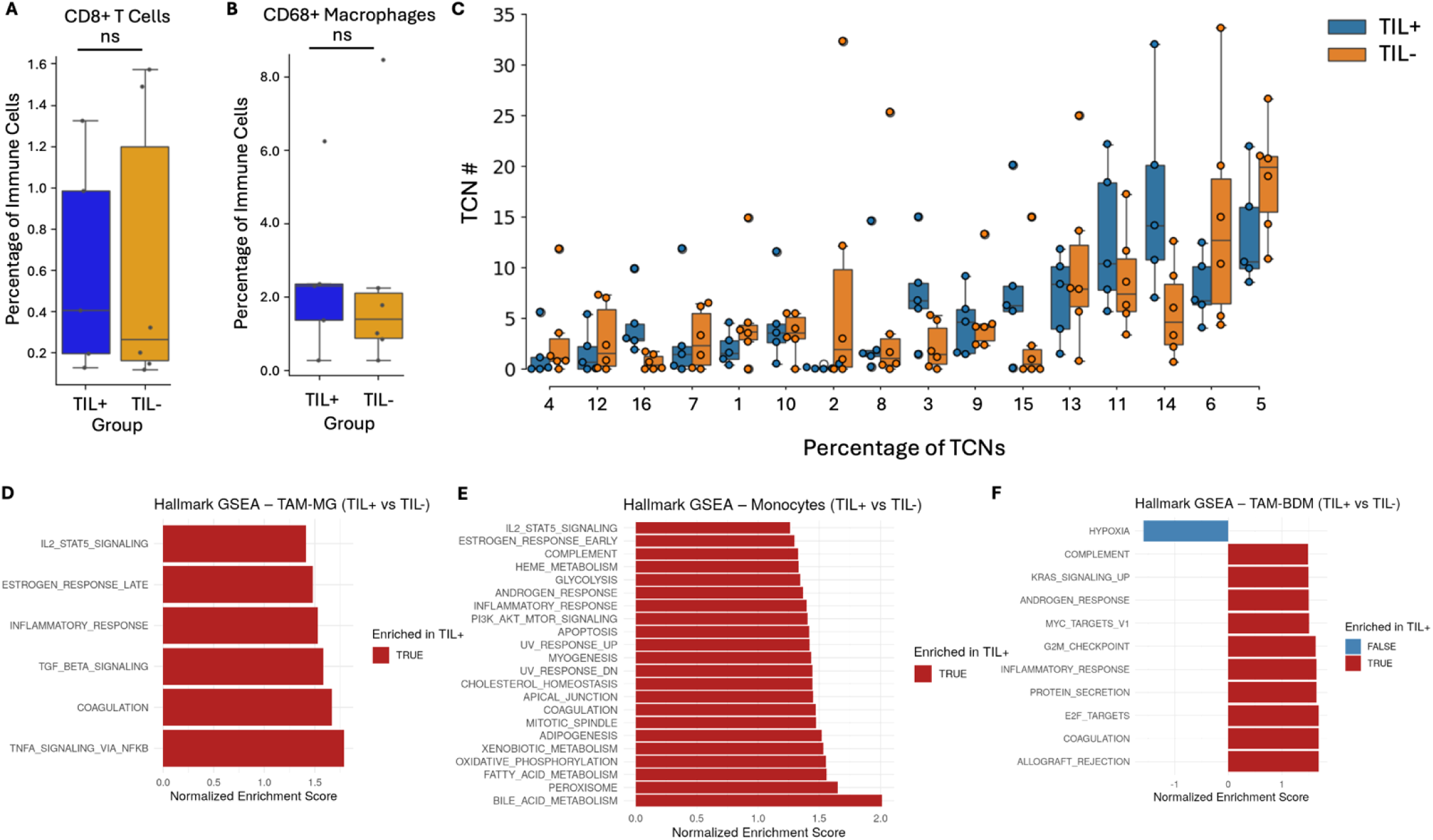
Myeloid transcriptional programs and conserved immune cell abundance according to TIL expansion status. A, Microglia-like TAM enrichment. Hallmark GSEA of microglia-like TAMs showing enrichment of inflammatory pathways in TIL⁺ tumors. B, Monocyte enrichment. Hallmark GSEA showing inflammatory and metabolically active programs in monocytes from TIL⁺ tumors. C, TAM-BDM enrichment. Hallmark GSEA showing inflammatory and antigen-presentation programs in TIL⁺ tumors and hypoxia-associated signaling in TIL⁻ tumors. D, CD8⁺ T-cell abundance. Similar proportions of CD8⁺ T cells across groups. E, CD68⁺ macrophage abundance. Similar proportions of CD68⁺ macrophages across groups. F, CD4⁺ T-cell-centric neighborhood abundance. Relative abundance of all T-cell-centric neighborhoods across groups. No significant differences were observed.

## Supplementary Tables

**Supplementary Table 1.** Clinical characteristics of the study cohort; Patients ranged from 45 to 79 years of age and received heterogeneous treatments, most commonly temozolomide, with additional use of bevacizumab, lomustine, and, in some cases, immunotherapy or investigational agents. Time to recurrence varied across the cohort, with two TIL⁻ cases showing no recurrence at last follow-up. Median time to progression was comparable between groups (TIL⁺ ∼234 days vs TIL⁻ ∼334 days). Overall survival ranged from 52 to 1203 days, with one patient alive at last follow-up, and showed no consistent difference between groups (median TIL⁺ ∼362 days vs TIL⁻ ∼408 days). No clear clinical or treatment-related features distinguished TIL⁺ from TIL⁻ tumors.

| ID | TIL | Age | Sex | Setting | Recurrence (d) | OS (d) | Treatment |
| --- | --- | --- | --- | --- | --- | --- | --- |
| 22-0189 | + | 45 | M | Primary | 211 | 299 | TMZ; Bev; CCNU |
| 23-0153 | + | 79 | F | Primary | 256 | 424 | TMZ; Bev |
| 23-0257 | + | 68 | F | Primary | 22 | 52 | None |
| 23-0227 | + | 63 | M | Recurrent | 758 | 1203 | TMZ; CCNU; Bev |
| 23-0196 | + | 51 | M | Primary |  |  |  |
| 23-0258 | - | 79 | M | Primary | NR | 142 | TMZ |
| 23-0080 | - | 49 | M | Primary | 417 | 632 | TMZ; CCNU; Bev |
| 24-0025 | - | 70 | F | Primary | 622 | Alive | TMZ; CCNU |
| 22-0225 | - | 51 | M | Primary | NR | 265 | None |
| 22-0160 | - | 54 | M | Recurrent | 251 | 780 | TMZ; Lerapolturev; pembro; Bev; CCNU |
| 23-0217 | - | 45 | F | Recurrent | 169 | 408 | TMZ; CCNU; pembro; Bev |

**Abbreviations**
TIL, tumor-infiltrating lymphocyte; OS, overall survival; d, days; NR, no recurrence; TMZ, temozolomide; Bev, bevacizumab; CCNU, lomustine; pembro, pembrolizumab; Lerapolturev, a recombinant poliovirus.

**Supplementary Table 2.**
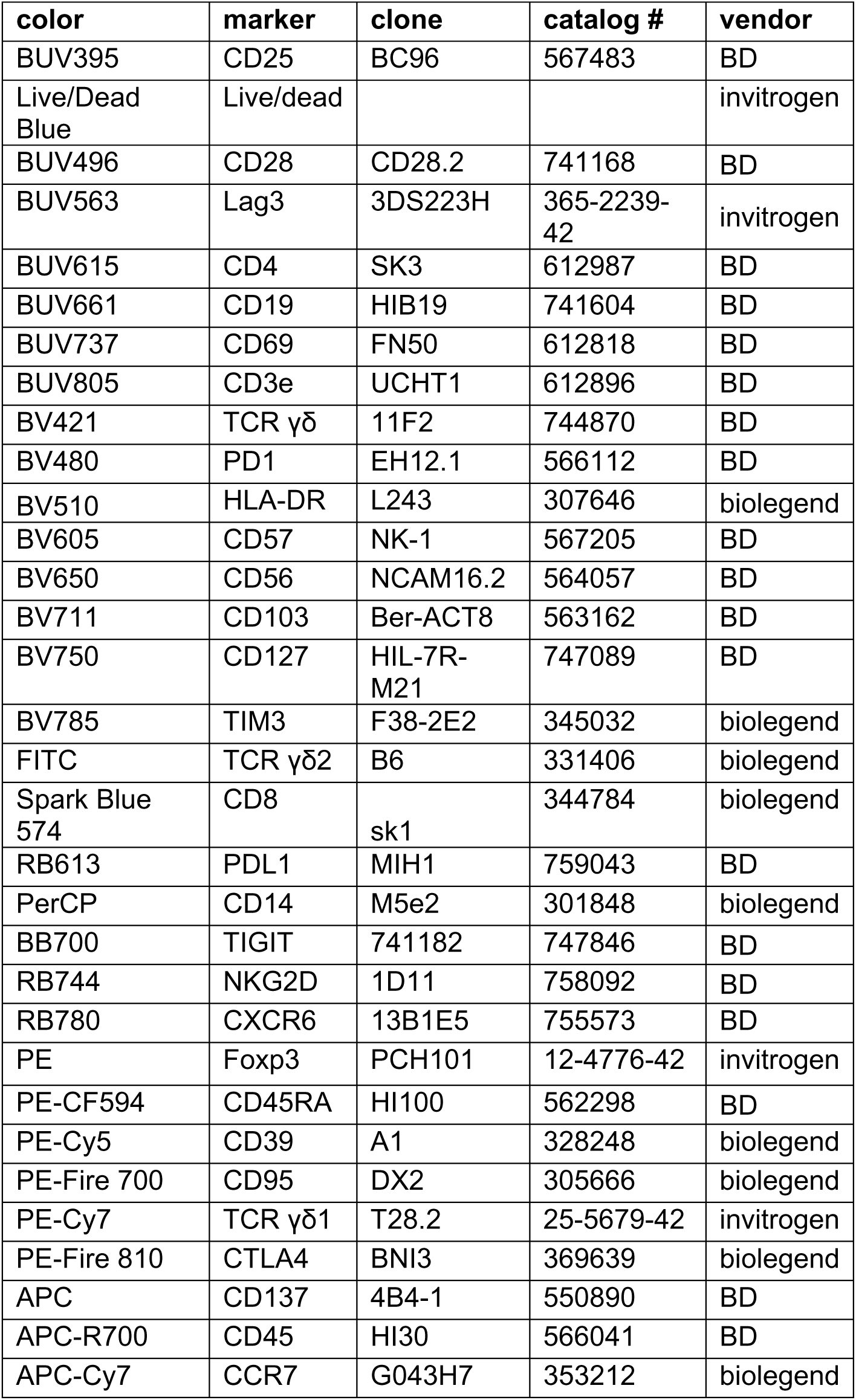
Antibodies used for TIL flow cytometry assays.

**Supplementary Table 3.** Antibodies used for the Co-Detection by Indexing (CODEX) assays.

| **Antibody** | **Clone** | **Company** | **Catalog Number** | **Oligo** |
| --- | --- | --- | --- | --- |
| Arg1 | Polyclonal |  |  | 43 |
| aSMA | Polyclonal | abcam | AB5694 | 69 |
| B-cat | 14 | CellMarque |  | 51 |
| BCL2 | 124 | CellMarque |  | 41 |
| CD11b | EPR1344 | abcam | ab209970 | 28 |
| CD138 | DL-101 | Biolegend | 352302 | 76 |
| CD163 | EDHu-1 | Novus | NB100-64980 | 45 |
| CD20 | rIGEL/773 | Noovus | NBP2-54591 | 48 |
| CD206 | E2L9N | CST | 49243SF | 55 |
| CD25 | 4C9 | CellMarque | custom | 24 |
| CD3 | MRQ-39 | CellMarque | MRQ-39 | 77 |
| CD31 | C31.3+C31.7+C31.10 | Novus | NBP2-47785 | 68 |
| CD34 | QBEnd/10 |  |  | 38 |
| CD36 | D8L9T |  |  | 74 |
| CD38 | E7Z8 | CST |  | 66 |
| CD4 | EPR6855 | abcam | ab181724 | 20 |
| CD45 | 2B11 + PD7/26 | Novus | NBP2-34287 | 56 |
| CD56 | MRQ-42 | CellMarque | custom | 29 |
| CD68 |  |  |  |  |
| CD73 | Polyclonal | BD | AF5795 | 75 |
| CD8 | C8/144B | Invitrogen | MA5-13473 | 8 |
| CD90 | EPR3132 |  |  | 21 |
| DCN | Polyclonal | Thermo |  | 63 |
| FOXP3 | 236A/E7 | Invitrogen | 14-4777-82 | 61 |
| HLADR | EPR3692 | abcam | ab209968 | 62 |
| Ki67 | B56 | BD | 556003 | 6 |
| MMP9 | L51/82 | Biolegend | 819701 | 80 |
| p16 | G175-405 |  |  | 59 |
| p53 | D07 |  |  | 52 |
| pH2AX | EP854(2)Y |  |  | 57 |
| Vimentin | RV202 | Novus | NBP1-97672 | 7 |

## Supplementary Information

Available upon request.

